# *Rp-vasa*: a *bona fide* Primordial Germ Cell marker that drives embryonic expression in the Chagas disease vector *Rhodnius prolixus*

**DOI:** 10.64898/2026.02.09.704890

**Authors:** G. Martins, M. Berni, T. Guedes-Silva, D. Bressan, J. Vieira, M. Cardoso, A. Pane, V. Gantz, E. Bier, H.M Araujo

## Abstract

*Rhodnius prolixus* is an insect vector of the protozoan *Trypanosoma cruzi*, the causative agent of debilitating Chagas disease, which is transmitted to humans during blood feeding. Identifying germline markers is a critical step in advancing vector control and transgenic technologies of these medically important insects. Transmission of genetic traits to the next generation requires proper differentiation of the germline that gives rise to gametes. Germline precursors are established during early stages of development as the primordial germ cell (PGC) population. Among the genes required for this process, *vasa* homologues exert a conserved role in germline specification. Here, we characterize and validate the genomic structure of the *R. prolixus Rp-vasa* locus and assess its expression during early embryogenesis. We observe widespread *Rp-vasa* expression in preblastoderm embryos. Later, during the cellular blastoderm and at the beginning of gastrulation, *Rp-vasa* and *Rp-piwi2* expression is restricted to PGCs, morphologically identifiable as a cluster of cells at the posterior of the embryo. We also report, for the first time, the use of *R. prolixus* regulatory sequences to drive the expression of exogenous genes. We identify the *Rp-vasa* regulatory region and show that these cis-regulatory sequences are sufficient to drive *Cas9* and *dsRed* expression in the early embryo. Together, these findings demonstrate that *Rp-vasa* has great potential for use as a PGC marker and as a driver for gene expression in transgenic and gene editing approaches for Triatomine vectors.

## Introduction

*Rhodnius prolixus* (*R. prolixus*) is a Chagas disease vector and model for molecular and genetic studies among triatomine bugs (Nunes-da-Fonseca et al., 2017). Infected kissing bugs deliver *Trypanosoma cruzi* (*T. cruzi*), the causative agent of Chagas disease, through their feces during blood feeding or by ingestion of food containing insect remnants (Chagas, 1909; Lidani et al., 2019). *R. prolixus* is an obligate hematophagous triatomine: both nymphs and adults feed exclusively on blood, and blood meals are tightly coupled to development and reproduction. In juvenile instars, a blood meal initiates the endocrine program that drives each molt. In adult females, blood feeding triggers the physiological and transcriptional cascade that promotes ovarian growth, vitellogenesis, and egg laying. Since blood meals create repeated opportunities to acquire *T. cruzi*, *R. prolixus* acts as a potential Chagas disease vector at several developmental stages (Lazzari et al., 2013). As vectors for a debilitating chronic disease that affects over more than six million people worldwide (Cucunuba et al., 2024; Monteiro et al., 2018; WHO, 2013), the development of molecular tools for biological control strategies in triatomines is paramount. With a sequenced genome (Mesquita et al., 2015), successful functional studies have been performed in *R. prolixus* using RNA interference (RNAi) strategies (Araujo et al., 2006; Berni et al., 2014; Brito et al., 2018; Lavore et al., 2012; Paim et al., 2013). However, no tools are yet available that enable the generation of precise loss-of-function mutants and transgenic lines (Araujo, 2025). In order to produce insect lines carrying heritable genetic modifications or transgenes, it is essential to identify promoter sequences for germline expression. In addition, genes that specify the germline could be used as targets for biological control by using them to engineer sterile insects.

The first embryonic cells to adopt a germline fate are known as primordial germ cells (PGCs) (Ewen-Campen et al., 2010; Extavour & Akam, 2003; Schwager et al., 2015). These cells exhibit highly heterochromatic nuclei, resulting in low transcriptional activity and rendering them unresponsive to transcription factors that drive somatic gene expression (Extavour & Akam, 2003; Strome & Lehmann, 2007). Two distinct modes of PGC specification have been identified: cytoplasmic inheritance (or preformation) and zygotic induction. Cytoplasmic inheritance has been extensively studied in the fruitfly *Drosophila melanogaster*. In *Drosophila*, a specialized cytoplasm called germ plasm, rich in maternal mRNAs and protein granules, is posteriorly localized in the oocyte and, during embryogenesis, incorporated into the posterior pole cells. Pole cells give rise to PGCs, become transcriptionally repressed and cellularize early, before the cellular blastoderm is formed (Chen et al., 2025). A functional germ plasm is required to form PGCs, and ectopic germ plasm is sufficient to induce a germ cell fate (Illmensee & Mahowald, 1974). Zygotic induction, on the other hand, requires signals from neighboring somatic cells, such as mesodermal cells, frequently long after gastrulation has taken place (Extavour & Akam, 2003). Within the hexapod clade, establishment of PGCs by inductive mechanisms is considered ancestral, with independent gain and loss of germ plasm seen in several groups during the evolutionary process (Quan & Lynch, 2016). While inductive mechanisms have not been addressed in kissing bugs, reports have failed to identify the presence of germ plasm in the hemiptera *Oncopeltus fasciatus* and *R. prolixus* (Ewen-Campen et al., 2013; Heming, 1994; Mellanby, 1935).

Regardless of the mode of PGC specification, the germline marker Vasa is a highly conserved DEAD-box RNA helicase that plays a central role in germline development across metazoans (reviewed in (Lasko, 2013). First identified in *Drosophila melanogaster* (Hay et al., 1988; Lasko & Ashburner, 1988; Schupbach & Wieschaus, 1986), *vasa* loss-of-function “grandchildless” mutants display a recessive maternal-effect sterile phenotype, producing viable embryos that lack PGCs. High levels of Dm-Vasa protein are found at the germ plasm during embryogenesis, where it facilitates the assembly of germ granules and translation of germline mRNAs (Dehghani & Lasko, 2015; Jeske et al., 2017). In addition, Vasa is crucial for maintaining genomic integrity in germ cells by promoting retrotransposon silencing via the piRNA pathway (Liang et al., 1994; Xiol et al., 2014). For instance, *Drosophila* Vasa is an essential Piwi-interacting RNA pathway component. Piwi proteins are a family of Argonaute proteins with roles in transposon silencing, germline development and the control of mRNA stability (Santos et al., 2023). *R. prolixus* harbors four *PIWI* genes: *Rp-piwi2*, *Rp-piwi3* and *Rp-ago3*, but not *Rp-piwi1*, are expressed in ovaries and are necessary for female adult fertility (Brito et al., 2018). Intriguingly, *Rp-piwi2* is highly expressed during early embryogenesis coinciding peak with a marked production of ∼28 nt long piRNAs, which suggests a critical role for this gene in the silencing of both resident and horizontally transmitted transposons (de Brito et al., 2024).

Homologues of *vasa* have been identified in a wide range of animals and are widely recognized as molecular markers of the germline in evolutionarily divergent clades. These include nematodes, mollusks and vertebrates (Fabioux et al., 2004; Fujiwara et al., 1994; Komiya et al., 1994; Roussell & Bennett, 1993; Tsunekawa et al., 2000; Yoon et al., 1997). Vasa homologues have been studied in arthropod models such as the spider *Parasteatoda tepidariorum* (Schwager et al., 2015), the mosquito *Anopheles gambiae* (Papathanos et al., 2009), the beetle *Tribolium castaneum* (Schroder, 2006), and the bug *Oncopeltus fasciatus* (Ewen-Campen et al., 2013). Importantly, *vasa* homologues serve as reliable germline markers in species that specify PGCs through either induction or preformation. This highlights the strong evolutionary conservation of *vasa* and its critical germline function, making it a particularly valuable candidate for investigation in triatomines.

In *R, prolixus*, PGCs have been identified based on cytological criteria. In early stages of embryogenesis, the PGCs are found at the posterior end of the germ band, where they divide asynchronously, before migrating into the yolk as the germband rudiment moves posterior-ventrally (Heming, 1994; Kelly, 1989; Mellanby, 1935). Molecular markers for these precursors that would enable in depth studies of germline specification have not been identified, however. Here we report on the identification and characterization of two putative PGC markers in *R. prolixus*. We find that *Rp-vasa* and *Rp-piwi2* are expressed during early embryogenesis, with highest levels in PGCs. We molecularly characterize both coding and regulatory sequences of the *Rp-vasa* locus. This analysis reveals that Vasa protein contains all conserved motifs of the DEAD-family helicases that are required to function in germline specification and maintenance. Finally, we identify *Rp-vasa* cis-regulatory sequences and show that they drive faithful germline expression during early embryogenesis. These studies provide an essential first step in the development of future transgenic and gene editing procedures.

## Methods

### Insect rearing

*R. prolixus* rearing was performed at 28°C (± 1°C) and 70-75% humidity, 12h light-dark cycle, with animals fed on rabbit blood throughout all five nymph stages until reaching adulthood at approximately 5-6 months. Animal care and experimental protocols were conducted following the protocol approved by the Ethics Committee on Animal Use at UFRJ, under registration 123/22. Technicians dedicated to the animal facility at the Institute of Medical Biochemistry (UFRJ) conducted all aspects related to rabbit husbandry under strict guidelines to ensure careful and consistent animal handling.

### *in silico* identification of germline markers and RACE PCR

*D. melanogaster* Vasa was used as a reference to search for *vasa* homologs in the *R. prolixus* genome using the BLASTp tool available on the VectorBase.org website. As in this initial version of the genome assembly only partial sequences were returned, to identify full length cDNAs we performed 5′ and 3′ RACE PCR using the FirstChoice® RLM-RACE Kit. The reactions were carried out according to the protocol. The resulting sequences were aligned using MUSCLE (Edgar, 2004). Primers are listed at Supp. Table 1. The identification and sequencing of genomic sequences between exons 5 and 6 was performed using converging and opposing primers designed based on the RproC3.5 version of the genome, followed by nested primers for Sanger sequencing. The resulting sequence was confirmed by Trinity analysis. Intronic primers are listed at Supp. Table 1.

*D. melanogaster* and *Anopheles sp.* Zpg, Exu, Ovo, Lola and Pum were used as reference to search for homologs in the *R. prolixus* genome using BLASTp as above.

### Cloning of the *Rp-vasa* promoter

After identifying the full-length hypothetical *Rp-vasa* cDNA, 4 kb upstream of the *Rp-vasa* initiation codon as well as 1 kb downstream of the hypothetical stop codon were cloned into pCR2.1-TOPO. To assemble the plasmid containing the *vasa* promoter driving *Cas9* and *dsRed* expression, a NEBuilder HiFi DNA Assembly reaction was performed according to the manufacturer’s protocol. Primers are listed at Supp. Table 1.

### *in situ* hybridization and immunofluorescence

Eggs were collected and embryos aged to the specified stages (Berni et al., 2014) at 28°C. Synchronized eggs were briefly washed with distilled water to remove debris and transferred to a 2 mL microtube with 1 mL of distilled water. The eggs were kept at 98°C for 90s, the water was replaced by 1 mL of formaldehyde 12% (in phosphate buffered saline, PBS) and fixed for 2 h (6-8°C). After this procedure, the embryos were incubated with 1 mL of formaldehyde 4% containing 0,1% of Tween 20 (PBST) under agitation for at least 1 hour. After fixation, the embryos were washed three times with PBST and transferred gradually to 100% ethanol for storage at −20°C.

For *in situ* hybridization (ISH) the embryo chorion was removed manually with fine forceps under a dissecting scope. Extreme care must be taken since PGCs are easily lost during manipulation. After dechorionation, embryos were re-fixed in 4% formaldehyde in PBST for 30 min at room temperature and, after a final wash with PBST, the embryos are gradually transferred to 100% ethanol and stored at −20°C or directly used for *in situ* hybridization with Digoxigenin-labeled antisense RNA probes following a previously reported protocol (Berni et al., 2023). Primer sets for probe synthesis are displayed in Supp. Table 1. Immunofluorescence in frozen sections was performed, following *in situ* hybridization, with primary anti-phosphotyrosine antibodies (1:1600; Cell Signaling) and secondary anti-mouse Alexa 488 (1:500, Molecular Probes). DAPI (1:500) was used to reveal nuclei. Whole mount ISH and sections were analyzed in a Leica SP5 confocal microscope.

### Frozen sections

ISH embryos were prepared for sectioning after defining their developmental stage as fixed whole mounts. Embryos were gradually transferred from PBST to OCT (Tissue-Tek, Sakura Finetek, Torrance, CA, USA), in steps of 2 hours each, starting with 25% OCT and reaching 100%. Once fully immersed in OCT, each embryo was positioned in a cut Eppendorf cap to orient its anteroposterior and dorsoventral axes and kept at −20°C until the OCT solidified. Sagittal 20μm sections were obtained with a semi-automatic cryostat Leica CM1860, at −18°C.

### Embryo injections

To overcome the hard, chemically impermeable chorion, 0-3h old embryos were lightly sanded at the dorsal–posterior end with #500 3M sandpaper. For this step, we used a custom-adapted 12V dental sander to ensure consistent abrasion. After 30 min - 2h desiccation in silica gel, embryos were aligned on double-sided tape and injected using borosilicate or aluminosilicate capillaries. 9 nL of purified plasmid DNA (1.3 ng) in injection buffer was delivered with a Nanoject II (Drummond Scientific Company) at a 45° angle to the filed site. The injection point was sealed with commercial cyanoacrylate (Loctite), and embryos were kept at 18°C for 16h and subsequently 28°C for 32h in a moist chamber.

### *in silico* analysis of regulatory elements

The 4kb *Rp-vasa* promoter sequence was defined by Sanger sequencing and analyzed using the MEME algorithm (Multiple EM for Motif Elicitation; meme-suite.org) to search for putative transcription factor binding sites. The 4kb was fragmented into 40 groups of 100 base pairs each, and 8 groups of 500 base pairs, in two rounds of MEME, to increase the likelihood of detecting repeated patterns characteristic of regulatory motifs. Subsequently, we used TOMTOM (Motif Comparison Tool) to compare each of the repeats detected with available online databases (https://jaspar.elixir.no/) for known motifs from *Drosophila*.

### Embryonic RT-PCR and qRT-PCR

In order to produce the cDNA library, total RNA was extracted from injected embryos or different embryonic stages as in Berni et al., 2014, or from ovaries dissected 7 days after blood feeding using Trizol Reagent (Invitrogen- cat: 15596026) as per manufacturer instructions. Total RNA was treated with RNAse free Turbo DNAse (Ambion, Life Technologies - cat: AM2238) to remove genomic DNA traces. cDNA was synthesized from 1ug total RNA using *in vitro* High-Capacity cDNA Reverse Transcription Kit (Applied Biosystems). Quantitative Real Time PCR (RT-qPCR) was performed on a QuantStudio 3 Real Time PCR system™ (Applied Biosystems) using PowerTrack SYBR Green Master Mix™ (Applied Biosystems). The relative gene expression was calculated using the ΔCT method (Livak & Schmittgen, 2001), using *Elongation factor 1* (*Elf1*) as reference gene. Oligonucleotides used for RT-qPCR were used in the final concentration of 150 nM and are listed in Table S1. All assays were conducted with biological triplicates and three to four technical replicates.

### Statistical analysis

All graphs and statistical analyses were performed using GraphPad 8 software (GraphPad Software, San Diego, CA, USA). Quantitative data obtained from qRT-PCR experiments were analyzed using one-way analysis of variance (ANOVA), followed by Dunnett’s multiple comparisons test. Choriogenic ovaries were used as the control group for all comparisons. Data are presented as mean ± standard error of the mean (SEM).The level of significance is shown in each Fig.ure (\*\*\**p* ≤ 0.001, \*\**p* ≤ 0.01, \**p* ≤ 0.05).

## Results

### Identification of putative PGC markers

We have previously identified several genes with sequence similarity to loci involved in the establishment of the *Drosophila melanogaster* germline such as *nanos*, *pumillio, piwi-2* and *vasa* (Brito et al., 2018; Coelho et al., 2021; Mesquita et al., 2015). Using antisense RNA probes, we detected *in situ* expression of *Rp-vasa* and *Rp-piwi2* in the tropharium and previtellogenic stage egg chambers during *R. prolixus* oogenesis (Brito et al., 2018). However, since chorionated eggs and embryos are impermeable to proteins and RNA, *in situ* analysis of gene expression has not been possible previously during this key developmental time period. We surmounted this technical barrier by developing a novel fixation protocol to enable *in situ* detection during the cleavage and blastoderm stages in which the chorion is removed and used this method to investigate *Rp-vasa* and *Rp-piwi2* expression during early embryogenesis. Whole mount *in situ* hybridization (ISH) shows that both *Rp-vasa* (Fig. 1A,B) and *Rp-piwi2* (Fig. 1C,D) are expressed in the posterior pole of the embryonic blastoderm, coinciding with the region where PGCs have been previously reported (Heming, 1994; Mellanby, 1935). After germband extension, both loci are expressed in the head region and close to the posterior growth zone, where PGCs reside above the mesoderm (Heming, 1994)(Fig. 1C,F). We also examined the expression of the homologs for *exuperantia* (*Rp-exu*) and *zero population growth* (*Rp-zpgA and Rp-zpgB*) (Supp. Fig. 1 and Table 1), fundamental genes for germline establishment in *Aedes aegypti* and *Anopheles stephensi* mosquitoes (Ellis et al., 2022; Hammond et al., 2021; Hughes, 2014). *Rp-exu* and *Rp-zpgA* displayed ubiquitous expression during embryogenesis (Supp. Fig. 1), while no expression was observed for *Rp-zpgB* during these stages (not shown). These broad expression patterns are unlikely to play a specific role in the establishment of PGCs. No expression for *Rp-nanos* (RPRC002927) has been reported during oogenesis, but this gene is expressed during 24-48h embryogenesis (around gastrulation and germ band extension), while *Rp-pumillo* and *Rp-staufen*, which display great similarity to *D. melanogaster* pole plasm genes, are expressed in unfertilized eggs and blastoderm stage embryos (Table 1) (Pascual & Rivera-Pomar, 2022). While the ubiquitous expression of both *Rp-exu* and *Rp-zpgA* does not support a direct function of these genes in the establishment of PGCs, their maternal expression might be linked to more general processes required also for germline specification, a question that would require further investigation.

**Figure 1.**
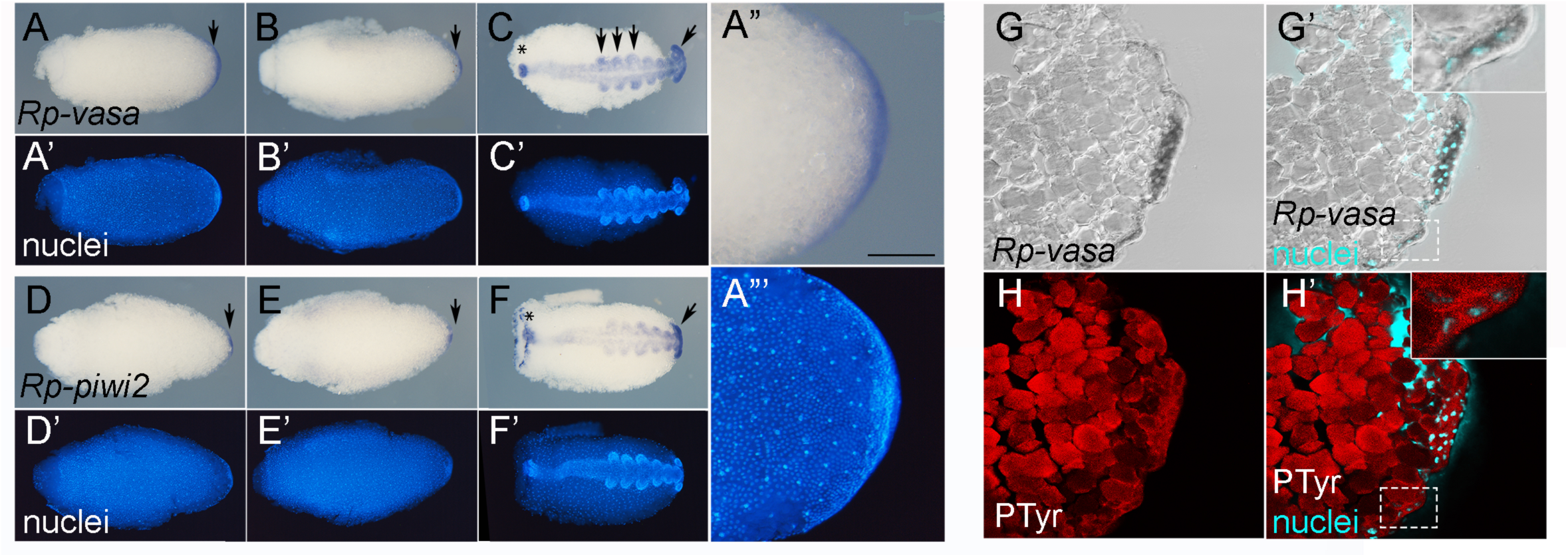
*Rp-vasa* and *Rp-piwi2* are expressed in Primordial Germ Cells. Gene expression pattern for *Rp-vasa* (A-C) and *Rp-piwi2* (D-F) defined by whole mount *in situ* hybridization (purple) and nuclei stained with DAPI (A’-H’). During the cellular blastoderm stage (stage 2; A,D) and early gastrula (stage 3A; B,E), expression is observed in posteriorly localized cells, seen more clearly in higher magnification (A”-A’”). After germband extension is complete (stage 4; C,F) both genes are expressed in the head (arrows) and the posterior growth zone (asterisk). *Rp-vasa* is also expressed in thoracic appendage buds (arrows in C). (G,H) Frozen sections of *Rp-vasa* whole mount ISH (G,G’) counterstained with anti-phosphotyrosine (H,H’) highlights the plasma membrane of cells that express *Rp-vasa*, confirming PGC expression at the blastoderm stage (stage 2). The central vitellum on the left displays high background fluorescence. Anterior is left, posterior to the right. Ventral views of the egg in (A,B,D,E,G,H) dorsal in (C,F).

**Table 1.** Expression of potential germline precursor markers. . Data derived from transcriptomic analyses from (Brito et al., 2018) (1), (Coelho et al., 2021) (2), and (de Brito et al., 2024; Pascual & Rivera-Pomar, 2022) (3). Previt., previtellogenic ovaries; Chor., choriogenic ovaries; Vitel., vitellogenic ovaries; Emb., embryo. Each study reports gene expression using different units (RPKM, CPM, or qualitative scoring). Therefore, expression levels are represented using a semi-quantitative scale based on the number of “+” symbols: + (>10 FPKM/CPM/score), ++ (>40), +++ (>100), ++++ (>400), and +++++ (>1000). NE (no expression) indicates genes that were not detected in the analyzed conditions or transcriptomes. * previously reported partial sequence for *Rp-vasa*.

| locus |  | Previt<sup>1</sup> | Previt<sup>2</sup> | Chor<sup>2</sup> | Vitel<sup>3</sup> | Emb<sup>3</sup> |
| --- | --- | --- | --- | --- | --- | --- |
| *vasa* * (C-term) | RPRC009661 | ++++ | +++ | +++ | NE | ++++ |
| *vasa* (N-term) | RPRC017810 | NE | NE | NE | NE | NE |
| *piwi-2* | RPRC002460 | ++++ | +++ | ++ | NE | +++++ |
| *nanos* | RPRC002927 | NE | NE | NE | NE | ++++ |
| *pumillo* | RPRC000720 | ++ | +++ | +++ | +++ | +++ |
| *staufen* | RPRC010019 | +++ | +++ | +++ | NE | +++++ |
| *exuperantia* | RPRC001415 | ++ | +++ | ++ | ++++ | ++++ |
| *zpgA* | RPRC014101 | +++ | +++ | ++ | ++++ | ++++ |
| *zpgB* | RPRC008769 | + | + | NE | ++++ | NE |
| *ovo* | RPRC002781 | +++ | +++ | +++ | NE | ++++ |

We sought to define the mechanism of PGC specification in *R. prolixus* by investigating in more detail the onset and pattern of *Rp-vasa* expression in the early embryo. We selected 0-6h and 6-12h old embryos, which represent preblastoderm and blastoderm stages of embryogenesis (Berni et al., 2014), respectively, to characterize more precisely the dynamics of *Rp-vasa* expression (Fig. 2). We observed ubiquitous *Rp-vasa* expression in the early blastoderm (Fig. 2A), that progressively becomes restricted in a posteriorly -ocalized pattern once the embryonic rudiment moves towards the ventral posterior region of the egg (Fig. 2B,C). At this stage (stage 2) the blastoderm is cellularized, as shown by anti-phosphotyrosine staining in frozen sections of *in situ* stained embryos (Fig. 1G,H). Cells expressing *Rp-vasa* are arranged as a loose group of internalized cells, covered by an epithelial layer of blastoderm cells, consistent with the arrangement previously described for PGCs (Fig. 1G,H) (Heming, 1994). We defined the location of the *Rp-vasa+* cells in relation to the prospective germ layers by performing ISH for the mesodermal marker *Rp-twist* (Fig. 2D,E). *Rp-twist* stains a large domain of ventral blastoderm cells that are displaced posteriorly with the posterior movement of the embryonic rudiment (Fig. 2D; Berni, Mota, 2023). Analysis of frozen sections revealed that this domain spans one cell diameter and covers the adjacent internalized PGCs at the beginning of gastrulation (stage 2B; Fig. 2E). The temporal quantitative analysis of *Rp-vasa* expression conforms to the *in situ* pattern shown in Fig. 2F.

**Figure 2.**
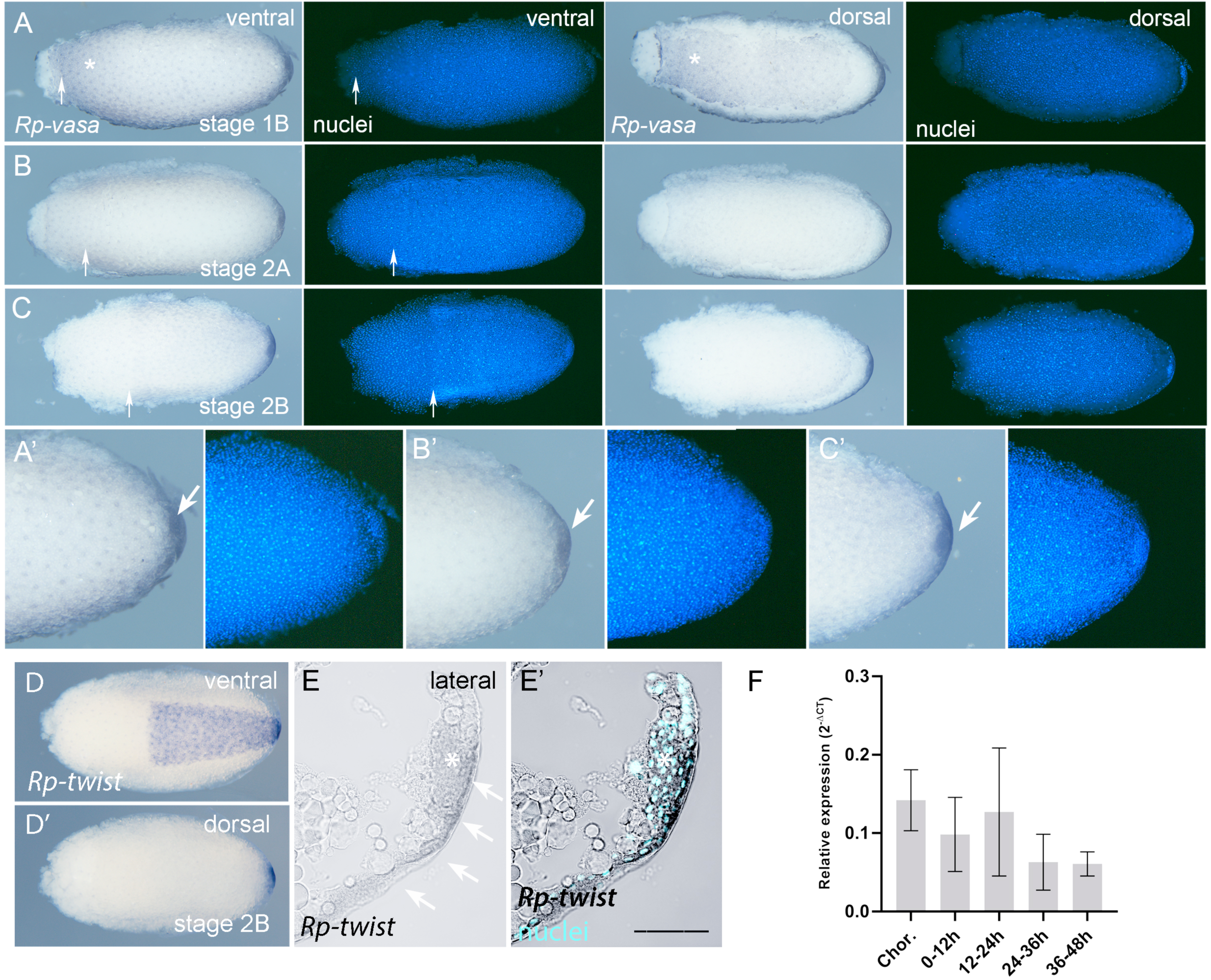
Widespread *Rp-vasa* expression becomes restricted to posterior PGCs by the end of the blastoderm stages. Whole mount *in situ* hybridization for *Rp-vasa* (purple) and nuclei stained with DAPI (blue fluorescence) shows widespread expression in ventral (A-C) and dorsal (A’-C’) regions of the early blastoderm (stage 2A; A-A’’), progressively becoming restricted to ventral and posterior regions in the late blastoderm (stage 2B; B-B’), and high levels restricted to PGCs shortly before and at gastrulation (stage 2B to 3A; C-C’’). (A’’-C’’) are high magnification of (A-C). The arrows in (A-C) point to the anterior position of the germ rudiment, that is displaced posteriorly with time. Asterisks in A,A’ show expression in the ventral (A) as well as dorsal (A’) blastoderm, the later completely lost in subsequent stages. (D) *in situ* hybridization for *Rp-twist* to reveal the localization of PGCs (arrows) in relation to the *Rp-twist+* prospective mesoderm in frozen sections of a stage 2B embryo. In A-D anterior is left, posterior to the right. Stages defined as in Berni et al, 2014. (E, E’) Frozen sections of *Rp-twist* whole mount ISH as in D. Shown are *in situ* only (E) and merge of *in situ* and nuclear stain (DAPI, E’). Arrows indicate *Rp-twist* staining along cells that compose the one-cell wide cellular blastoderm, asterisk denotes internalized PGCs. (F) Quantitative RT-PCR shows changing levels of *Rp-vasa* expression during early development. Data are presented as mean ± standard error of the mean (SEM). No statistically significant differences were detected by one-way ANOVA with Dunnett’s multiple comparisons test, using choriogenic ovaries as the control.

### Structure of *Rp-vasa* genetic locus and encoded Vasa protein

Given the germline expression of *Rp-vasa* in *R. prolixus* PGCs, the widespread use of *vasa* homologues as PGC markers, and the successful use of *vasa* promoters for germline expression in *D. melanogaster*, *A. gambiae* and *C. quinquefasciatus*, we experimentally tested potential regulatory sequences of the *Rp-vasa* locus. A previous study identified the partial sequence of *Rp-vasa* (annotated as RPRC009661 in the current RproC3 version of the *R prolixus* genome) that lacked the initiating codon as well as 5’ sequences where the *Rp-vasa* promoter most likely resides. We performed Rapid Amplification of cDNA Ends (RACE PCR, see methods) to acquire the complete mRNA sequence and identify the N-terminus of protein coding sequences (Fig. 3A; Supp Table 1). Extending the *Rp-vasa* sequence allowed us to identify the full Dead and Helicase C domains of the predicted Vasa protein (Fig. 3B), which display great sequence conservation when compared to a broad spectrum of insect species (Supp. Fig. 2).

**Figure 3.**
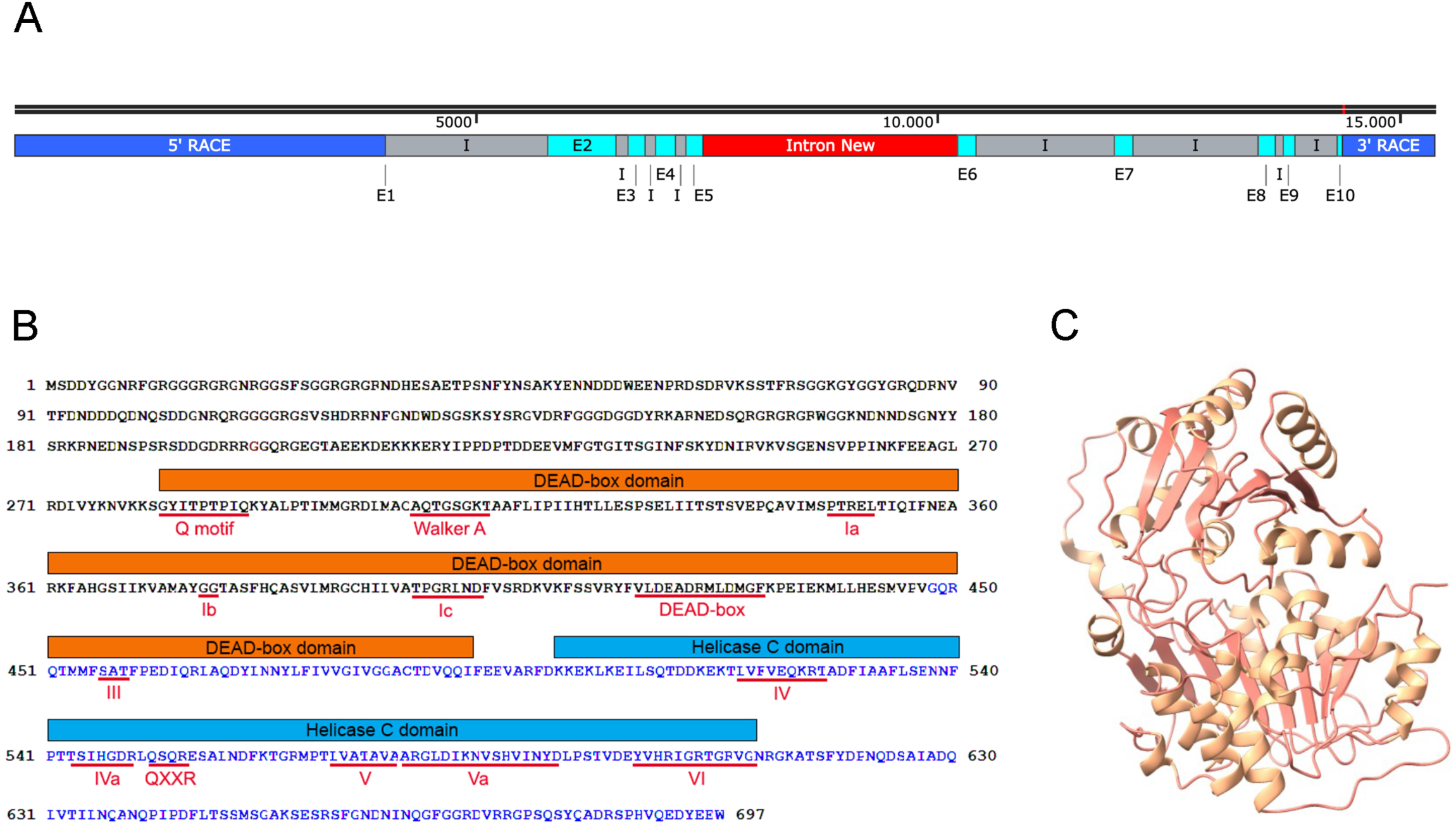
Structure of the *Rp-vasa* locus. (A) The 5’-UTR, coding sequences and 3’-UTR of *Rp-vasa* are distributed among two different sequences in Vectorbase (green and yellow). RACE analysis and Sanger sequencing shows that they are transcribed in the same direction. The complete coding sequence spans 10 exons (blue). Sequencing of an uncharacterized intron (red) using primers from exons 5 and 6 (Fig.ure S3) confirms exon arrangement and reveals the complete structure of the *Rp-vasa* locus. (B) Rp-Vasa protein displays all the conserved motifs characteristic of insect Vasa. In black is displayed the sequence previously reported as RPRC017810 and in blue the sequence previously reported as encoded by RPRC009661. Conserved domains of Vasa proteins are highlighted. (C) Rp-Vasa protein structure predicted using Alphafold. Shown are amino acid residues 235-690 containing conserved motifs.

We next aligned the experimentally determined cDNA sequence against the *R. prolixus* genome and recovered two predicted homologous gene sequences: RPRC017810 and RPRC009661, which align respectively to the N-terminal and to the C-terminal sequences of our predicted *Rp-vasa* open reading frame. The current annotation places these genes 17567bp apart, transcribed in opposing directions and from different DNA strands, with an intervening region composed of highly repeated sequences. We explored the possible continuity of these sequences as part of the *Rp-vasa* locus by designing PCR primers to amplify and sequence the intervening genomic region that our cDNA sequence had predicted there should be between exons 5 (part of current RPRC017810) and 6 (part of RPRC009661) of *Rp-vasa* (Fig. 3). Indeed, Sanger sequencing of this amplified product plus Trinity analysis of RNA-Seq sequences that had not previously been allocated to the genome assembly (Grabherr et al., 2011) confirmed the continuity predicted for exons 5 and 6. Therefore, as a result of our RACE and exon5/6 breakpoint sequence analysis we propose a gene structure for *Rp-vasa* in which the currently annotated RPRC017810 or RPRC009661 sequences are inverted and are in actuality comprise a single locus. In this corrected gene structure, *Rp-vasa* is composed of 10 exons and 9 intervening introns (Fig. 3A; Supp. Fig. 3), similar to the gene structure reported for the hemiptera *Cimex lenticularis vasa* locus (LOC106660862).

The predicted protein sequence is also in agreement with a complete Vasa homolog. Proteins of this family present 11 conserved motifs, namely: Walker A, Motif Ia, Motif Ib, Motif Ic, Walker B (DEAD-box, or Motif II), Motif III, Motif IV, Motif IVa, Motif V, Motif Va and Motif VI. These are part of two domains: DEAD-box (or DEXDc), containing motifs Q to III, and Helicase domain C (HELICc), with motifs IV to VI (Fig. 3B). Vasa proteins are distinct from other DEAD-box helicases by having an additional Q Motif at the N terminal of Walker A, and, between Motifs IV and V, they have a conserved Q--A motif (Adashev et al., 2023; Banroques et al., 2011; Gustafson & Wessel, 2010; Linder & Jankowsky, 2011; Tanner et al., 2003). Motifs Q, Walker A, DEAD, and VI bind to the ATP molecule, while motifs Ia, Ib, IV, and the first portion of motif V interact with target RNAs. Meanwhile, motif III and the final portion of motif V connect the ATP and RNA interaction sites (Banroques et al., 2008; Linder & Jankowsky, 2011; Schroder, 2006; Xu et al., 2021). The 3D structure of Rp-Vasa predicted by AlphaFold is also in agreement with that determined for *Drosophila* Vasa by X-ray crystallography (Fig. 3C) (Sengoku et al., 2006). In addition to the conserved motifs, Rp-Vasa displays large Intrinsically Disordered Regions (IDRs) at the N- and C-terminus. IDRs have been shown to facilitate intermolecular interactions that lead to phase separation/transition, resulting in the formation of condensates of biomolecules such as RNPs (Xu et al., 2021). Interestingly, the Dm-Vasa N-terminal IDR is required for pole plasm localization (Liang et al., 1994).

### Regulatory sequences in *Rp-vasa* drive embryonic expression

We searched for regulatory sequences that may drive expression of *Rp-vasa* in PGCs by scanning for putative regulatory elements 4kb upstream of the initiation codon. MEME analysis identified predicted sites for binding by four different transcription factors, based on regulatory elements described for *D. melanogaster*: Ovo, Lola, Sna and Eve (Fig. 4A). Among the four, Ovo and Lola putative binding elements presented the highest similarity to *Drosophila* sequences. *ovo* encodes a transcription factor with Zn-finger domains that acts as an evolutionarily conserved component to regulate germline development (Hayashi et al., 2017). *Lola* encodes a transcription factor that plays a role in *Drosophila* oogenesis and spermatogenesis (Bass et al., 2007; Davies et al., 2013). A great number of sequences with similarity to *lola* were identified in our analysis. Identifying the precise *lola* ortholog will be an interesting goal for future studies. In the case of *ovo*, only a single candidate ortholog was identified (RPRC002781) (Lavore et al., 2015; Pascual & Rivera-Pomar, 2022). This locus has also previously been reported as *Rp-shavenbaby* (*svb*) (Tobias-Santos, 2019). Based on the conserved germline function shown for *ovo* orthologs, we hereafter refer to this gene as *Rp-ovo*. *Rp-ovo* expression (Table 1 and Fig.ure 4), is consistent with maternal (Pascual & Rivera-Pomar, 2022) and zygotic functions. Parental RNAi knock-down of *Rp-ovo* results in loss of embryonic viability and in embryos with a poorly extended germband (Tobias-Santos, 2019). The breadth of expression and the knock-down phenotypes suggest that *Rp-ovo* performs multiple functions during oogenesis and embryogenesis, which may include an evolutionarily conserved function in germline establishment. Although no *eve* (*even skipped*) ortholog has been identified in the *R. prolixus* genome, *Rp-sna* (RPRC005707) expression has been previously reported as a presumptive mesoderm marker in the early embryo (Berni et al., 2014; Mesquita et al., 2015). It will be interesting to investigate whether *Rp-ovo* and *Rp-sna* exert an effect on *Rp-vasa* expression by future functional analysis using RNAi. Based on the uniform distribution of putative regulatory elements along a 4 kb region, upstream of *Rp-vasa*, we were unable to define a regulatory subdomain responsible for early embryonic gene expression. Hereon, we refer to this 4Kbp region as the *Rp-vasa* promoter.

**Figure 4.**
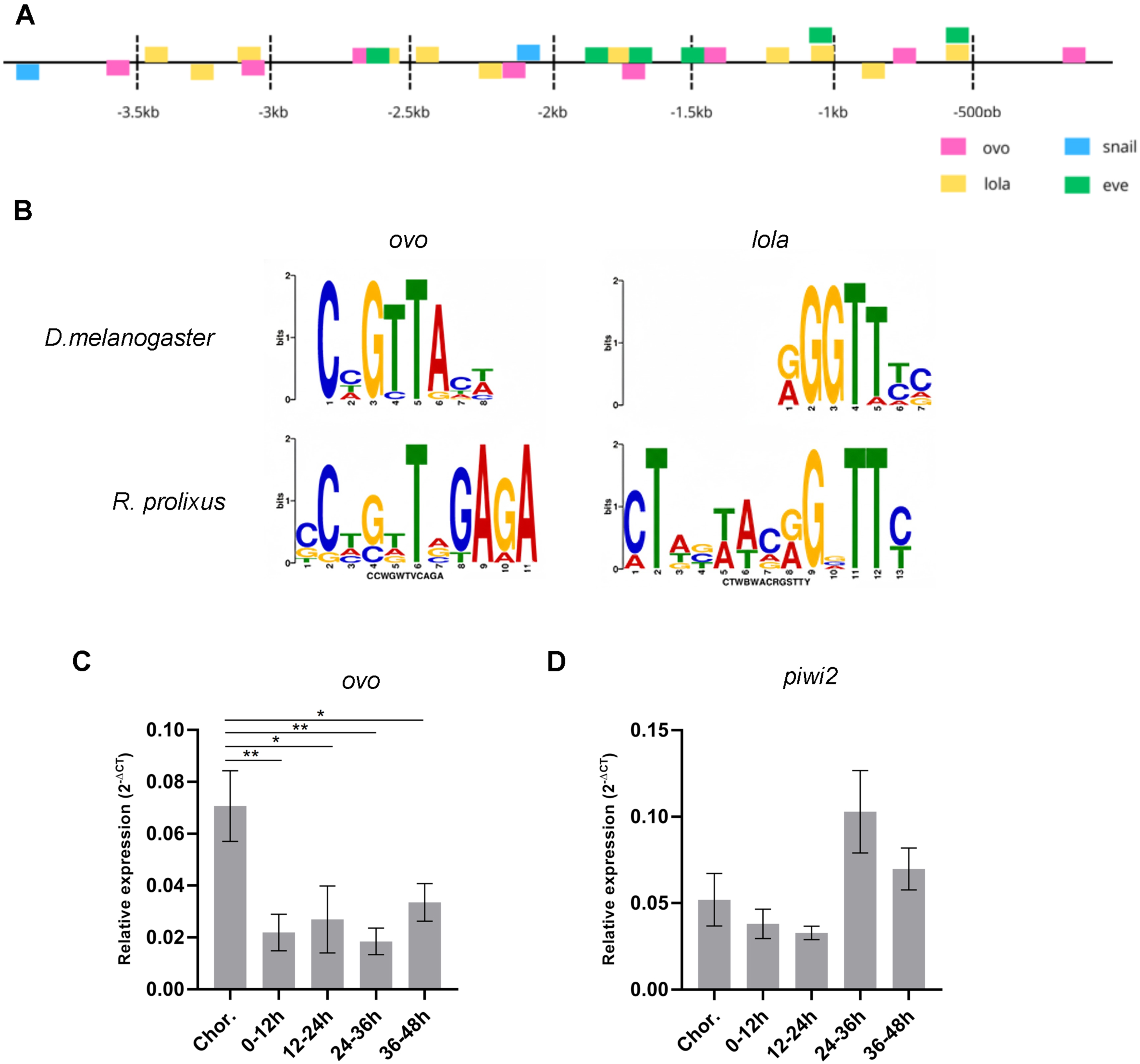
Identification of putative regulatory sites in the *Rp-vasa* promoter. (A) MEME analysis identifies putative binding sites for several transcription factors, distributed along the entire 4kb intergenic sequence upstream of RPRC017810. (B) Sequence similarity between previously characterized *Drosophila* Ovo and Lola binding elements and putative Rp-Ovo and Rp-Lola elements identified *in silico*. (C, D) Expression of *Rp-ovo* (C) and *Rp-piwi2* (D) during early embryogenesis defined by qRT-PCR. Data are presented as the mean ± standard error of the mean (SEM). Statistical analysis was performed using one-way ANOVA followed by Dunnett’s multiple comparisons test, with choriogenic ovaries used as the control.

To test whether this 4kb sequence could drive faithful reporter gene expression, we constructed a vector with this *Rp-vasa* promoter sequence, driving expression of the nuclease Cas9 and the fluorescent marker DsRed, followed by 3’ sequences from *Rp-vasa* that we defined by RACE. We injected purified plasmid DNA into pre-blastoderm stage embryos (0-3h old; stage 1), aged them for 48h (reaching late blastoderm and early gastrulation under the low temperature condition), and then extracted RNA from these embryos for qRT-PCR and sequence analysis. Indeed, *Cas9* and *DsRed* mRNA were observed only in embryos injected with the vector but not in the control (Fig. 5B). We confirmed that the amplified sequences were generated from RNA and not from contamination by the plasmid by using primers against the plasmid backbone (Pl) (Fig. 5B). In the future, it will be interesting to further delineate minimal enhancer elements within the 4kb sequence that are required for PGC expression.

**Figure 5.**
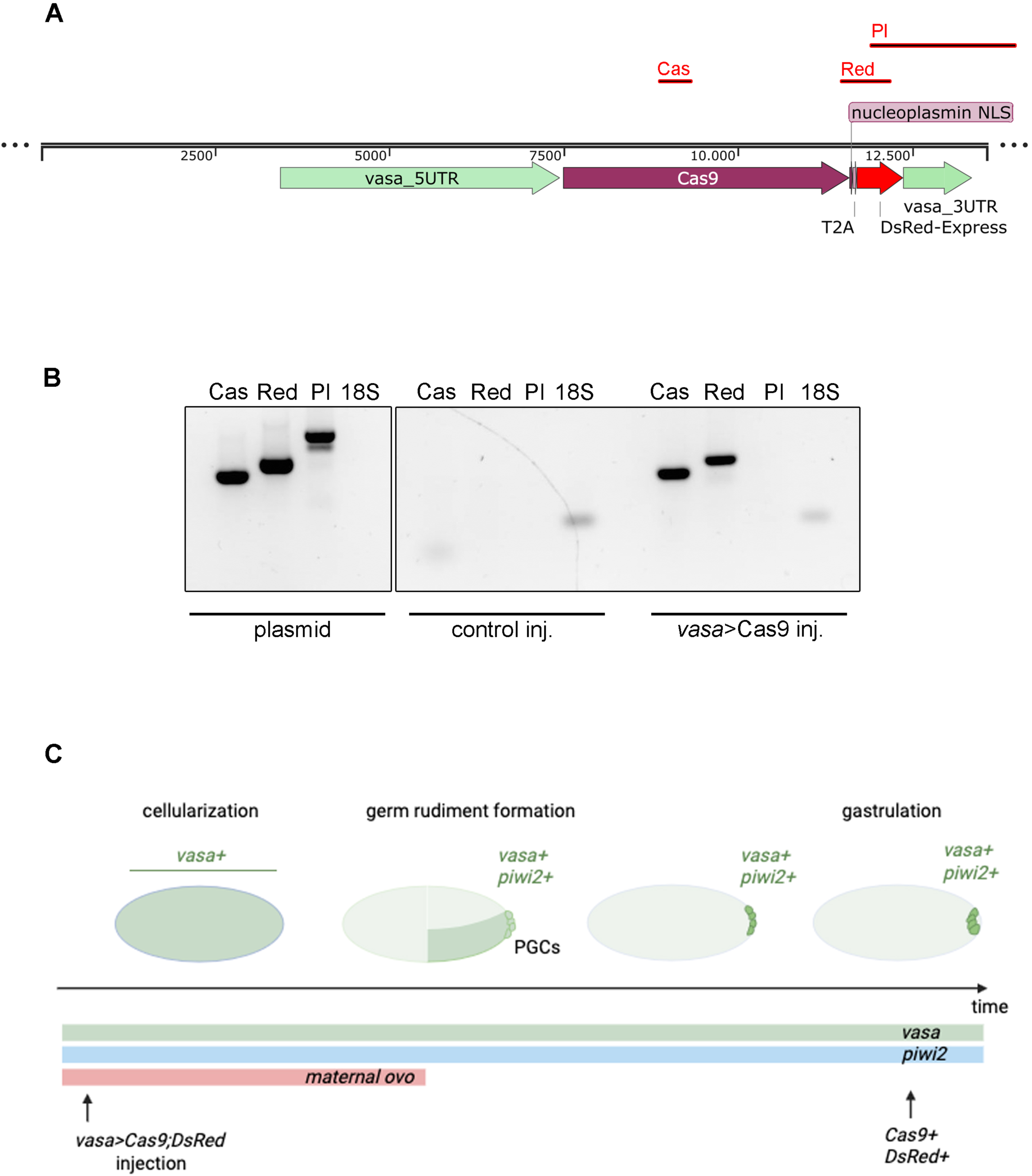
The *Rp-vasa* promoter directs gene expression during early embryogenesis. (A) Features of the *vasa>Cas9;dsRed* vector (B) Expression of *Cas9* and *DsRed* under the control of the *Rp-vasa* promoter is detected around gastrulation (stages 2B to 3A), in embryos that had been injected with the *Rp-vasa*>*Cas9; DsRed* vector during the cleavage stage (stage 1). (C) Schematics summarizing the expression of PGC markers *in situ* and the temporal expression of putative germline regulators during early *R.prolixus* embryogenesis.

## Discussion

### Molecular identification of PGCs in *R. prolixus*

Previous cytological analysis identified *R. prolixus* PGCs presence at the posterior region of the embryonic rudiment, prior to gastrulation (Heming, 1994; Mellanby, 1935). Based on the conserved function of *vasa* and *piwi* homologues in germline establishment (Ku & Lin, 2014; Yajima et al., 2014), the high expression levels of *Rp-vasa* and *Rp-piwi2* we observed in these posterior cells supports their identification as PGCs. Analyzing the full expression pattern from oogenesis to embryogenesis should help resolve their potential function in germline establishment. During oogenesis, *Rp-vasa* and *Rp-piwi2* are expressed in oocytes and trophic cells of the tropharium (Brito et al., 2018), suggesting that they are transcribed during oogenesis and transported to vitellogenic oocytes by trophic cords. The ubiquitous distribution of *Rp-vasa* transcripts identified in early embryos (stages 1A and 1B) is consistent with such maternal provision. This distribution may represent a conserved pattern in Hemiptera, since it was also described for *Oncopeltus fasciatus vasa* (Ewen-Campen et al., 2013). In *Gryllus*, in which PGCs are specified during late embryogenesis by mesodermal induction, *Gb-piwi* and *Gb-vasa* are ubiquitously expressed in the early embryo, displaying no signs of asymmetry (Ewen-Campen et al., 2013). This ubiquitous *Gb-vasa* expression is unlikely related to PGC specification. In *Drosophila*, which has a pole-plasm-dependent mode of PGC specification, *vasa* is expressed during oogenesis, and *vasa* mRNA is detected throughout the embryo at low levels during early cleavage stages (Hay et al., 1990). This early expression pattern may be related to the control of mitotic divisions, as reported for the fly and the sea urchin (Pek & Kai, 2011; Yajima & Wessel, 2011). Taking into account this evolutionary conservation, ubiquitous *Rp-vasa* expression may be related to a cell cycle function as well.

Later during the cellular blastoderm and early gastrula phases of development, we observed high levels of *Rp-vasa* and *Rp-piwi2* that are restricted to posteriorly localized PGCs. A similar distribution has been shown for *O. fasciatus vasa* (Ewen-Campen et al., 2013). Different from *R. prolixus*, however, *Of*-*piwi* expression is not observed in posterior *O. faciatus* PGCs (Ewen-Campen et al., 2013). In *Drosophila,* Vasa protein becomes asymmetrically localized to posterior cells of the syncytial blastoderm, restricted by the action of pole plasm assembly components such as *oskar*, *staufen*, *tudor* and *valois* (Hay et al., 1990; Lasko & Ashburner, 1988; Liang et al., 1994). The posterior localization of *Rp-vasa* and *Rp-piwi2* may be key to a function in PGC specification since in *Drosophila vasa* regulates the translation of germline mRNAs (Carrera et al., 2000) and *piwi* is also involved in germline determination (Megosh et al., 2006), conferring genomic stability as an essential player in the piRNA-mediated silencing of transposons (Teixeira et al., 2017). Accordingly, loss of *Dm*-*vasa* or *Dm-piwi* function results in the formation of fewer PGCs (Carrera et al., 2000; Teixeira et al., 2017). On the other hand, knockdown for *Oncopeltus Of-vasa* and the potential *vasa* regulator *Of-tudor* resulted neither in loss of germline precursors or decreased fertility, indicating that alternative or additional loci are required for PGC specification in this hemipteran. Parental knockdown for *Rp-piwi2* disrupts oogenesis by causing loss of trophocytes, egg-chamber degeneration and oogenesis arrest (Brito et al., 2018), unfortunately preventing functional analysis of a potential role in embryonic PGC specification by parental RNAi. Although we did not conduct functional assays of *Rp-vasa* and *Rp-piwi2* during embryogenesis—due to low survival rates of dsRNA-injected embryos and the extended time to sexual maturity (6 months) required for fertility assays—their early expression indicates a role in germline establishment. Furthermore, their regulatory regions could be useful for driving germline expression in transgenic studies (see below) and for identifying other factors involved in PGC regulation.

Contrary to *Rp-vasa* and *Rp-piwi2*, which display germline specific components of expression, *Rp-nanos*, *Rp-zpgA*, *Rp-zpgB* and *Rp-exu* are not promising as PGC markers or to drive germline precursor expression since these genes are either widely expressed (*Rp-zpgA*, *Rp-zpgB*) or not expressed in the germline (*Rp-nanos, Rp-exu*). *zero population growth* (*zpg)* encodes a gap junction innexin, which plays a crucial role in early germ cell differentiation and survival and has been shown to be required for germ cell development in *Drosophila* and mosquitoes (Tazuke et al., 2002; Thailayil et al., 2011). The mosquito *zpg* promoter has been demonstrated to express in a germline-specific manner (Hammond et al., 2021). However, the absence of specific expression in embryonic PGCs precludes the use of innexins, *Rp-exu* and *Rp-nanos* as *R. prolixus* PGC markers.

### Establishment of germline progenitors in the kissing bug

Given that *Rp-vasa* is a good marker for PGCs, it should be possible to find, among its regulatory regions, clues as to how expression in these cells is controlled, pointing to possible mechanisms of PGC specification. Despite the lack of evidence of a pole plasm in *R. prolixus* (Heming, 1994), the identification of PGCs early during development (pre-gastrulation) argues against a zygotically induced mechanism. However, immediately before gastrulation *Rp-vasa+/Rp-piwi2+* PGCs are located adjacent to the *Rp-twist+* ventral blastoderm cells. These cells will form the mesodermal germ layer after gastrulation. Our *in silico* analysis also identified two putative binding sites for Snail, a transcription factor that in *Drosophila* contributes to the development of the mesoderm and regulates the epithelial-mesenchymal transition in several developmental contexts (Ashraf & Ip, 2001; Leptin, 1991). While knockdown for *Rp-twi* and/or *Rp-sna* may reveal an inductive role of these genes in the prospective mesoderm for PGC specification, evidence of PGC induction by the mesoderm performed in other species has confirmed such as role only during later stages, after elongation of the germband (>stage 5 in *R. prolixus*).

The presence of Ovo binding elements upstream of *Rp-vasa* and the expression of *Rp-ovo* during oogenesis and early embryogenesis suggests an early function in PGC specification that should be addressed in future functional studies. *ovo* exerts a conserved function in germline establishment, despite the mode of PGC specification. For instance, *ovo* accumulates in PGCs and is required for *vasa* expression in *Drosophila* (Yatsu et al., 2008). Additionally, knockdown of mouse *Ovol2*, which is expressed in PGCs, decreases PGC number (Hayashi et al., 2017). It is noteworthy that different *ovo* isoforms display opposite actions during *Drosophila* development, calling for careful ontogenetic examination of *Rp-ovo* transcripts (Andrews et al., 2000). Nonetheless, it is important to point out that *R. prolixus* regulatory elements may display great sequence divergence in relation to *D. melanogaster* elements and thus might not be detected by our current analysis. Furthermore, the putative elements herein described should be tested by chromatin-IP and knockdown assays before they are assigned as actual regulatory sequences.

The presence of all conserved functional domains in the predicted Rp-Vasa protein is revealing about its putative function in early embryogenesis and, possibly, establishment of germline precursors. The approximately 400 amino acid central domain characteristic of DEAD-box helicases shows high identity to other insect Vasa (Supp Fig. 2). *Drosophila vasa* mutants for DEAD-box residues fail to restore the ability to form pole cells (PGCs) in *vasa-* embryos (Dehghani & Lasko, 2015), and DEAD-box mutants lose their helicase functions, resulting in loss of features such as pole cell specification and repression of transposon-encoded mRNAs. Additionally, the Vasa C-terminus contains a stretch of highly acidic residues (691-697 in Rp-Vasa), conserved only in Vasa homologs, that is required for *Drosophila* germ cell formation (Dehghani & Lasko, 2015). On the other hand, the large N-terminal IDR regions of Vasa proteins have been associated with interactions with ribonucleoparticles (Xu et al., 2021). The predicted AlphaFold 3D structure conserves the central core of the protein required for activity, indicating that Rp-Vasa displays all characteristics required to function as an active RNA helicase. Importantly, the predicted full-length protein sequence will be fundamental for the development of Rp-Vasa antibodies for protein and RNA co-immunoprecipitation studies, leading to the identification of additional players for establishment of the germline.

### Molecular tools for targeting kissing bug vectors of *T. cruzi*

Targeting insect reproduction with sterile insect techniques (SITs) has proven efficient for controlling and eradicating insect pests. Generation of sterile insects can be achieved through classical methods such as irradiation (Lees et al., 2015) or by the use of transgenesis (Burt, 2014) and self-propagating gene drives based on the CRISPR/Cas technology (Hammond et al., 2016; Kyrou et al., 2018). The molecular identification of PGCs in *R. prolixus* is a necessary first step to enable the development of these modern technologies in a vector of Chagas disease. First, *Rp-vasa* and *Rp-piwi2* could be used as markers for the effects of gene knockdown or knockout on the establishment of germ cells by accessing *Rp-vasa* and *Rp-piwi2* expression levels in the embryo by ISH or qRT-PCR, rather than by accessing fertility as adults. These early indicators are invaluable in species that take 5-6 months to reach sexual maturity such as the kissing bugs. Further, *Rp-vasa* and *Rp-piwi2* may prove useful in engineering sterile insect techniques (SITs) in kissing bugs. Their expression in the tropharium during oogenesis (Brito et al., 2018) and in the embryo PGCs (this report) indicates the potential to interfere with germline establishment at two independent moments, increasing the chances of generating sterile insects. Finally, studying genes that alter *Rp-vasa* and *Rp-piwi2* function may lead to the identification of additional targets for SITs.

Regulatory sequences that drive germline expression are essential for molecular techniques requiring the stable maintenance of transgenes or alleles across generations. Regulatory sequences of *D. melanogaster vasa* and *nanos* have been used to drive germline expression of Gal4 (Brand & Perrimon, 1993) and Cas9 (Sebo et al., 2014), among others. Similar drivers have been developed for *Anopheles* and *Culex* mosquitoes and the moth *Bombyx mori* (Carballar-Lejarazu et al., 2020; Feng et al., 2021; Gantz & Bier, 2015; Harvey-Samuel et al., 2023; Papathanos et al., 2009; Smidler et al., 2024). Here we have identified a regulatory region from the *Rp-vasa* locus that can be used to drive early embryonic expression. This 4kb sequence drives *Cas9* expression in stage 2-3 embryos, a period characterized by high levels of *Rp-vasa* expression in PGCs, and could be used in the future to generate transgenic bugs expressing *Cas9* in the germline to expand the use of gene editing tools in *Rhodnius* (Lima et al., 2024). The identification of putative Ovo and Lola binding elements along the promoter is in agreement with germline expression and should help determine how *Rp-vasa* expression is controlled. Additional promoter analysis should define the minimal sequences required for PGC expression in order to build a minimal driver. In short, the identification and cloning of the *Rp-vasa* promoter should enable the development of essential tools for transgenesis and gene editing in the kissing bug.

## Acknowledgements

We would like to thank members of the Araujo lab for helpful discussions and comments on the manuscript. We are grateful to the animal facility at the Institute of Medical Biochemistry for technical assistance with Rhodnius husbandry.

