## Supplemental material for "*Rp-vasa*: a *bona fide* Primordial Germ Cell marker that drives embryonic expression in the Chagas disease vector *Rhodnius prolixus*"

### Supplementary Tables and Figure legends

### Table S1. Primers used in this study

| name | forward | reverse | amplicon size (bp) |
|---|---|---|---|
| **ISH probes** | | | |
| *Rp-twist* | ggccgcggCCAGAACTAGTGCCTCTGTCG | cccggggcACCCTCCATTCTCCACACTG | 405 |
| *Rp-vasa* | ggccgcggCATTGTTGGAGGCGCTTGTA | cccggggcATGTAGCTTTGCCACGATTACC | 412 |
| *Rp-piwi2* | ggccgcggTACTGATTGGACATTATACCA G | ggccgcggTATCTGATCTCTGTGAAACATC | 519 |
| *T7 universal* | | AGGGATCCTAATACGACTCACTATAGG Gcccggggc | |
| **qRT-PCR** | | | |
| *Rp-vasa* | ATCGTGGCAAAGCTACATCC | GTCAGGTATTGGCTGATTTGC | 101 |
| *Rp-ovo* | GTCGCCGACTAGTTGTTCAC | GGCGTTGCAGGCTAAATGAT | 239 |
| *Rp-piwi2* | TGACTTCTATAAGGCAACACG | TATCTGATCTCTGTGAAACATC | 117 |
| *Elf1* | GATTCCACTGAACCGCCTTA | GCCGGGTTATATCCGATTTT | 92 |
| **cloning** | | | |
| Vasa promoter | CCTCGTTCGTAACCACTACACC | CATAATTGAAATTTGAATGCTACTTATTA ACAATTCTCTGGTAAC | 4011 |
| Vasa 3' UTR | TGAAAAGTGGAATTGATAAATAGATTAG CTCTGAATGCG | GCTATCCAACGTTAAGTTGTGAATTTAG TAGTGC | 1002 |
| Vasa promoter + Topo bckb Hifi | TTCTTCTTTGGCATAATTGAAATTTGAAT GCTACTTATTAACAATTCTCTGG | CTAAATTCACAAGCTCTTCCGCTTCCTC GC | 7674 |
| Vasa 3' UTR Hifi | CACCTGTTCCTGTGAAAAGTGGAATTG ATAAATAGATTAGCTCTGAATGC | AGCGGAAGAGCTTGTGAATTTAGTAGT GCACGACTGGC | 1009 |
| Cas9 + dsRED Hifi | ATTTCAATTATGCCAAAGAAGAAGCGG AAGGTCG | TCCACTTTTCACAGGAACAGGTGGTGG CG | 4949 |
| **sequencing** | | | |
| Vasa RACE 1179 | CCCCACCTACCACGCCCACGACCTCTT TGACTATC | | |
| Vasa 9660 For | GCAGAGGTGTGGACAGATTTGGTGGTG G | | |
| Vasa 9661 For | CTAGAAAGTTCGCCCATGGCTCC | | |
| Vasa 1120 For | CCACTCCTGGCCGACTAAATGAC | | |
| 1-2 | CCACTCCTGGCCGACTAAATGAC | CGCCTCCAACAATGCCCACAAC | 3000 |
| 3-4 | TTCCATTGTCGGGTCAAGAT | CCGGGTAATACAATTATCCATTC | 1800 |
| 5-6 | GGTAATTTTGGGAGAGTTGCGT | TCAAAATTAAAGGTGCTAGGCCA | 550 |

**Figure S1. Ubiquitous expression of *Rp-exu* and *Rp-zpgA* indicates no specific role in PGC establishment.** Ubiquitous gene expression is seen for *Rp-zpgA* (A,B) and *Rp-exu* (C,D), as defined by whole mount *in situ* hybridization (A-D; purple) and nuclei stained with DAPI (A'-D'). No specific pattern is observed for either gene during the blastoderm (stage 2) and germband extension (stage 3) stages.

**Figure S2. Complete Rp-Vasa protein sequence with conserved motifs.** Predicted *Rhodnius prolixus* Vasa protein sequence with conserved Vasa domains, based on MUSCLE alignment to Vasa from *Drosophila melanogaster* (NM_001273529.2), *Apis mellifera* (NP_001035345.1), *Cimex lenticularis* (XM_014383864.2), *Halimorpha halys* (XM_014423399.1) and *Gryllus bimaculatus* (AB378065.10). Motifs Q to DEAD box are part of the DEAD domain, while motifs III to VI are part of the Helicase C domain. Sequences previously reported as encoded by RPRC017810 end at amino acid position 447, while RPRC009661 starts at 448. Genomic and cDNA sequencing support the prediction that RPRC017810 and RPRC009661 are contíguous genes transcribed in the same direction, both part of the *Rp-vasa* locus.

**Figure S3. An intronic sequence between exons 5 and 6 corrects the *Rp-vasa* gene structure.** (A) DNA sequence amplification with inverted primers 1 and 2 located in exons 5 and 6 generates a 3kb fragment corresponding to an unpredicted intron. Nested primers 3 and 5 generate a predicted 1.8Kb band inside this new intron. Amplified products were used for Sanger sequencing to generate the full *Rp-vasa* genomic sequence presented in Figure S4. (B) Location of the primers used in relation to the proposed *Rp-vasa* genomic locus.

**Figure S4. Complete sequence of the *Rp-vasa* locus.**

**Figure S5. *R. prolixus* innexins aligned to insect orthologs.** Predicted *Rhodnius prolixus* Innexins (ZpgA and ZpgB) aligned to insect homologs.

**Figure S6. *R. prolixus* exuperantia aligned to insect orthologs.** Predicted *Rhodnius prolixus* Exuperantia protein aligned to insect homologs.

Martins et al. Figure S1

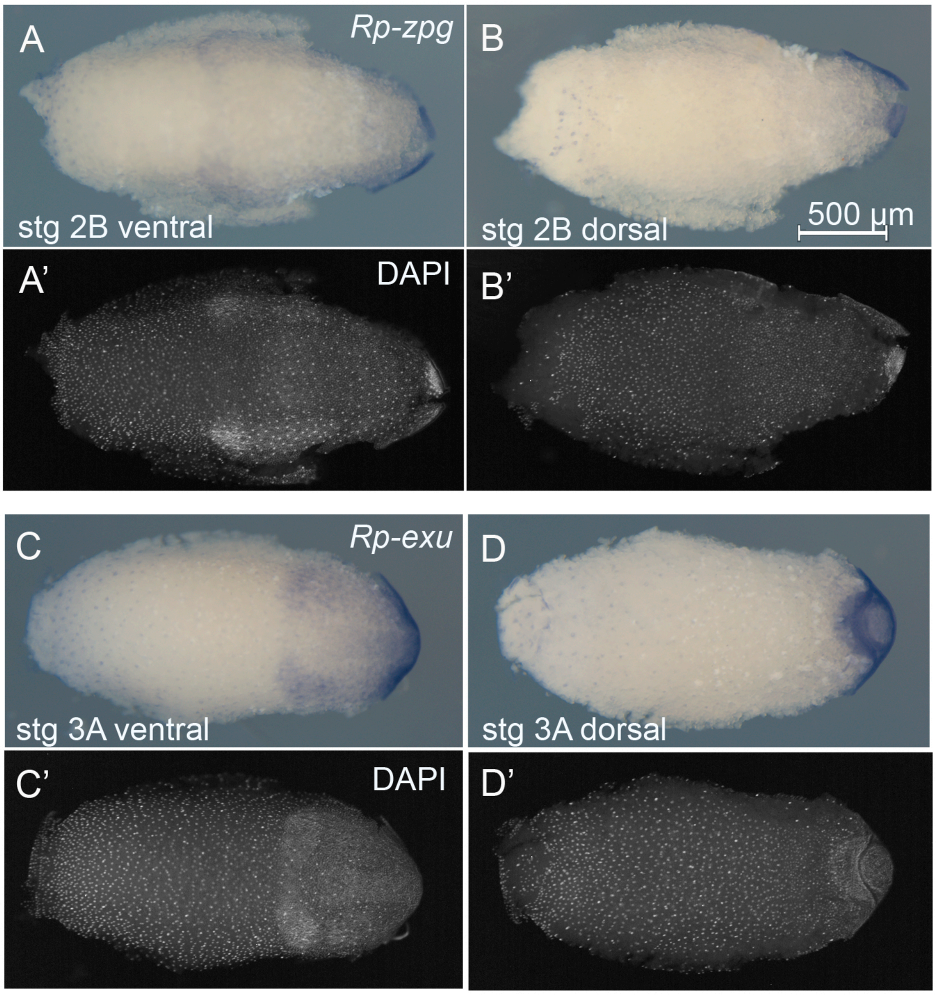

Martins et al. Figure S2

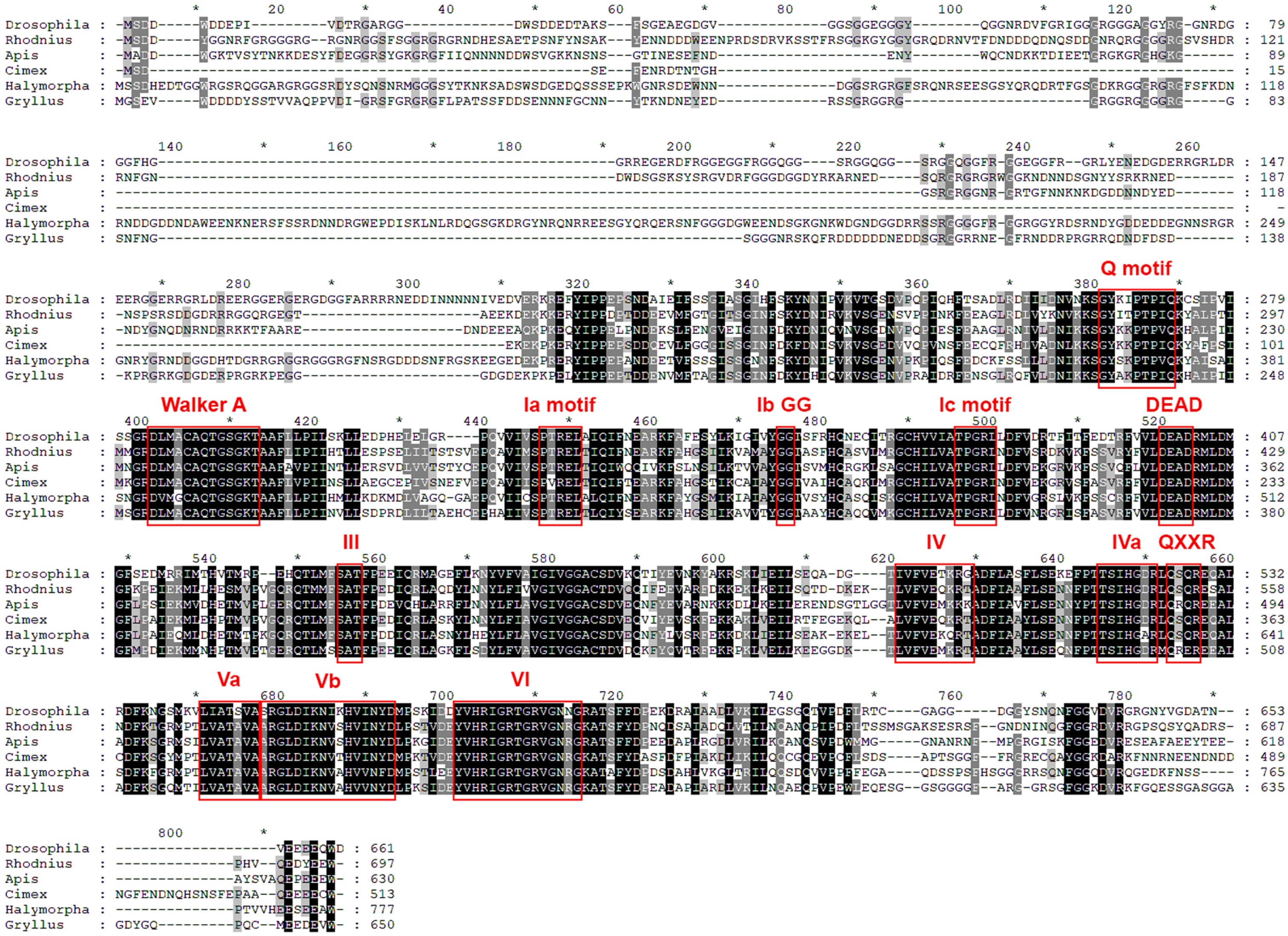

Martins et al. Figure S3.

A

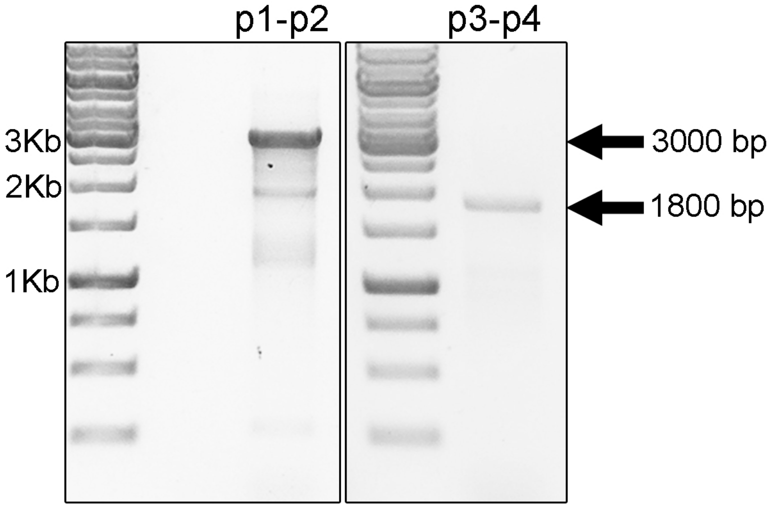


B

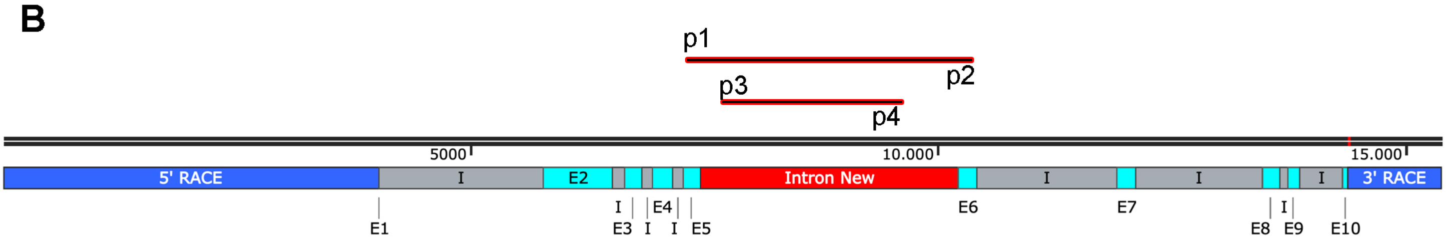

Martins et al. Figure S4

>5' mapped  (4011 bp)

CCTCGTTCGTAACCACTACACCACCCACCCCTAGCCCTAAGAATATCATGAACTATTCTAAGTAAAAAAAAAAAAATTCAT

TTGATACATTTCATCATAAAAAAAAAATATGAATTTGGTATTTTGAGGAGTCATCGGAATTAAGTCTAATTAAACTTGAGC

TAAAAATACATTTTACGTTTACACTTGGACGTAAACCGAAAATAACATGACTTAGGACAATTGTGCCAAGGTGAATTCAC

ACATATAAATATGAAAACAAAACAAATGATTTGCATTTGATTTTGAGACAAGTTTTGAAAATTTTTATAAACCATCTTTC

TTCGAAAATCGAATATACGGTATTTTCAGAGGAACAAATTAGAACATCTTGGTTAAACATTTTAAATCTGTAAGTAACAT

TTAACTGAAGGCAAGTTACAAAAATGAAAATAAATCTTGGGAAAATTCTCACCTAGCTCTCCTATTTCCATTTGTTCTTC

TGCATCTGGTGTCATGTTGCCATAAACTATTTCGGCTTCAGATAATTTATGCTTTTTAATGTGCTGCTGGTTCAAATATC

TGAAATAAGAGGTGAAAATAGTTCTTAACTTATATTTATTTATTTTTTATCACACTGGGAATACAATTTTTAGACATTAC

TAAATTTTAAATAGAAAATAGTTTCAAAATACTATTCTAACAAAATCCACTATTCTTACAGATAAAGCCGATGTAGATAA

TTTATTCCTTGGCTGTATTCATTCCAGGCATTATAATAATTTTTTAATAAGTTACTTTCCCCAGCAGAGACAACCTGAAA

CGACCATAAATTAAGCGAAATAGTACTTCTCTGTTCTGGAAATGATATTCATTAATGAAACATTAGTGTTGAAAAAAAAT

CTAACCTTTTCTAATAGATTTTTCACATGGTTAATCAGAAAGCGTCGGGTTTCTTGATACAAACGATCTGCAAGAGGTTC

TGGGTAAGCCACACACAACGAGTACACATCACTATAAGGGGGAAAAATAATAATATTAGATGACCAGAATCAGTTGACAG

ATGCAAATAATATGGAGTTAGGATAGAAGTTTTAAACAAGTTAAGCTATTTTACACATAGTAGCATAACATACACACATA

ATTCATTAAACACATGGAAAACCAATGAAAAATTAAAGCAAGCTAAAAATTGATAATTTAGTTTAGATTCGACCGGGACC

CATCAAGAAGCTTTAGGTCAAAACCTACATATAAACAAAATAAATAAATAAAGGACGCTTTACAAAATAGATTTTCCTGT

TACAGATTTTTCTTGAAAGTGACCTTCTTCAAACTTTGTATAACTTACTACACACATACTTGAAAAGTAGTAGAAACACT

ACTAGGGCTTTTCTCAGTTTCAGTTTAGAATCTTGTACGGTTTTAATAAAAAATTAACTATGCAACCTAATTTCAAAATT

TTTAGAAAATCTTTAAAGTTGAAGTCAACTTTACTGTTTTTTAATATCAAAAATGGTTCCCATGATTTACTTCAAACTTT

GAATACACATTAGATAGAAATATGTAGTCGTATTGAACTTAAACGTTCTTGCAGGAACTTAGCTTCAAGGAGGTTTTACA

GTAAAGTGGAGTCAAGCTTTCCTCTAATGTTCGATCTAAGCGAATTTTATGGGTCTAAGTAATATTTATAACTATCAAAC

GTGTTCAGAGTTTAAAAAAAAAAAAAAACTGAGTGGAAGCTCGATTATATTAAACATCTTTTTTTCTCGAATTTTGTTTGA

TTTTTTTACAAGATAATAAAGATAACCCAAATTATATGAACTTGAACCATATATTATTAGGCACAAAAGAATCTTTTCCA

GAAAGTGTGTAACTGCATTTCTGTACGGGCTGAGACGTCTACTTCCTCTATTTCGTTTTGTTTGTTGTAGCATGCAACAA

TTATTTTCAATTACTTCTTCAATTCTAATTCTTAATTTTTAGAAATCCTAATTACTTAGGCAAAAGTGAGAAAATAGTTT

GTTGTTTTCGCCATAAGAGTAGCCATTTTTACTATTTCTTACACTTTCATAACAATAACCAACAACTTTGCTCGAATGAG

ACATTTGAGAAAAAAAACAAAAAAAAATAGGGGCGCTGATTTTGAAGAGGATGCGGTAAAAGATAAAAAAAAACAAGTAA

CTTTCTAATCTAATCAGGGGATGAGGGCAAAGAGAAAAAATAACCAACTTTCTAATCTAATAATTAGTTATTTCACAAAT

GTGAACGATCTCTAGAACGGGTTCAAAACATTTTTTGGCACAAAACTCAGAATTAAATTGCTTGAGTCATGCTAACACGA

ACATCATTGCACATTAAATTTTAGAAATCTCATATAGAAAATGAGTCATTCACTAAATTGCGTTATAATCTAAAAGGAAA

AAAAAGTTTAGTTATCCTTGATTCAGCTATTTATAATATGACCTAGATAATATCGGACAGATTTACTCATATTGAATTTA

ATTATGTTAACAGGGAATTTCCTGATCCAAACAACTATATGTTACAAAAATAAACGCCAAATATTCTGTATATTTCTACA

GTCACATACATAAATAATAAATTGAAAATAAAATTCTTAGAAGAAAACGAATTTTAAAACTGTCTTGTAACCTTGTAAAG

GTTCTTTTCTTTGTTTTGTATACTATTTGTGACAAGGGGTAGAAGTCCTCTGCAAGACAAACATGAAATTTGGCACAGAC
AAATTATCTATAATTTCTTCTTCAATTTAAACCTTTCTCAACTATTGTAACAATAATAAAATCAAAAGTAAGTTGAGCAT
AATGATTTTTTTTAATTAGGCAAAATCAAATAGACCAAGAAAATAAAAAAAATCTGACTACCTGAATCGATCATTCCATGT
AGATCGTGGGACATTACCAAGTGTAATAACACCTTTAACAGTTTCTTGTAGTACGTCCCATGTTTCATCAAAATCTACTT
TCCTTGGTTTTAATGACATTTTTTATTCCTACAAAGATATGTCATACTATCTAAAAACATACACAAAATAGTTTTTTTTA
TTCCGCAAAAATCCAAAAAACCAGAAGAAAAATAACAGCTTGAGAATAAAATGATTGTAAATAATCGTTTTCAAATAACT
ACCATAAGTATATATGAAACCCGAAATATAATTTTAAAAAACTATAGTTACCATGAGTGGTCGTTTACTTGTAAAAAAAA
TAAAACTAAAAGATTTAAAACGAATTGATATTCCTCTATTTTCCTTTGAGGTGCAGAATGAACACGCACAGAACGTAATA
TTCGTGAATGACTTTACACCTTAGCTATCAACTTTAGTGACTATTTTCGACTCAGTTATACTTTCAGCTCTACTTTTTGT
CACCTCACTTTACTTCCAAACAGCTAAGCTAGCAAGCCATCATACTACAGGTTCAGCAATGTAGTAATAAAAAACTAATT
TCCCCATTTAATTTAATTTTTAATTAAAAATAATAATAATTAGTTATCTCCAACCTAAGACTTTTCGGTACCTTACAATT
AATAGCAACTTGTAAACATTTATTCCTTGAAGTATGAAATTTACTAATTTCTGAAGTAGCATTTTTTGTTAGTAAACAGT
TTTTATAAATCCTTATTACATTATTACATTAAGATTTTCTAAATTATTATCCTTTTATAAAAATTTAAAAAAATGCTAGA
AAATCAACATTAGATTAAAAACTTTTGTATTTATTTTGTGTAATATAATTTACTAAAGGCGTTTGAAAAAATCTTTACTT
TTTGGTAACTAATAATTTAATTAATAGTTTTTTTATTAACAAAAGTTGGAAACTAAAGATGTTATAAATCTCCCCTCCAC
ATGTTAAAAAAGGCTCCATAAATGCTGTTACAGAATACAATGGAGTTAGCGGTTGCCATTTTAACATTTTAAGTTTATCA
CGGTAAGCACGTGTGAAACTGAATTCGGAATTAGAACTTGTCTTATGTTACCAGAGAATTGTTAATAAGTAGCATTCAAA
TTTCAATTATG

>5' UTR VectorBase  (120 bp)

GCGGTTGCCATTTTAACATTTTAAGTTTATCACGGTAAGCACGTGTGAAACTGAATTCGGAATTAGAACTTGTCTTATGT
TACCAGAGAATTGTTAATAAGTAGCATTCAAATTTCAATT

>Exon 1  (7 bp)

ATGTCAG

>Intron 1 VectorBase  (1756 bp)

GTATTTGAACATTTCTTTTTTTAAATAAAATTATACACAGCATATCTTGTTACTTCTGGACATGTAAACCAGTACCGGCT
TTGTGGTTTCAGTGATAAATGAACCATGTTTTCACTTATTTAATTATTTTTTATTTTTTCAAGGCAGATAGAACTAATAA
AAGGAATATAGAAATTATTGTATGATTTCTTACGGTTTTTGGGGTTCTTAGCTGTGAAGAGAAACATAAATTTGATTGCG
TTAACAAACGATTTGTGTATGTAATAGAAAGAAAAAGTTACGTTCAGGTCTCTCTGTGTTTGCTATTTTTTAAGGAGTCC
TTGGCTTGAAGTTGGCTATGTATGGTATTTCGTAAAGAAATTTTGGTTCCGAAATTATGTCTCAGAAATAATAATACCGT
CTTATAAATAAGTGTCATTCCTCAGAGATTTATTGTTACCCGATATTTGGCATCCCAGGCTAAGAAGCTACCTCATTCCA
CTAATCCACAATTGTAACAATAATTATGCTTCATGCCTTTGCGTGGTGAAAACTATATTAGTACAGCAACTTACGAGAAA

TTATTTGTTATCTGATCTTAAATTTAGTGTTTGGTGGTATACTTAGGCGTAACATTTTAGAGAAATTAGTGACTTGTTAA
CTATTTGATTCCCAAGAGGTTAACAATGCCACATTTTTTAAATATCAAATTTCAGTAAAACTTGTTCGTATCTTAATTTT
TTATGAGATGCAGTAAAAGCTACCTAAGATTCTTAAAATTAAGAAATTTAATAAATTCTGTTTCTGCAGAAGAAATTTTC
AGAATGGCTCTTATTGAGCATTTTTTTTTTACTAAAAACTGATTAATTTGAATATATGATTAAATTACTTTAAAAAAAAT
AAATGATACGGGTTTGTTTTCTATTTATTTAGCAAAGTTTCAAATTGTTTTTGCTCAAGGTAGACATAGCAGTCAAAATG
TATGATTACTGAAAACGGGCTTTTAAACTTAACTATAATTAACTTCGAAATGGGTTGTTTGAATAACAAGTTTACTTTTC
CTCTAATGAGAACTCTACGTCTATATAAATCATCAAATGTTTCCAGTTCTTCTCTATTACATATTCTTGAAACCATTCTA
GCACATTTCTTTTTGTTGGAATGTTGGTAAGTTCAGTTTTTCAACATTTATTTTGTTGGTGTCACGAGTCAACATAAATT
GTACTCTCCCAGCAATCTATTGTATTTTCTTTGCCGCTCTGACGAGGACTATTTATTTTTTACTCTTACTTTTTTTGCTA
TTCCCCGTTTAAGTTAATGATCCTTACGCGTTTATACAGTATTTTATACTGTCATTGAAGATATTGTATCCAGTGAACAT
TAGATTAGGTATGGAATTACTTTCTTAACTGTTTTAACCCTTGGTCGTTGAGGAATTTCCATGATGTGTCAGATATATTC
TGCGATAAGTGAGAATCAATGACTTCTAGTGCAGGATTATATCAATAGGAACGATTATTTTGGTTTCTTTCATACCACTT
TCTGTTGCGCCTATTGTTGTTGACTTTTCTGAAGTTCACTTTTTAATATAATAATTTAAATAAGAATGAATAATAAAAAA
TTTCCATACAGTTTGGTGACTTTAATTAATTTGTGAACCTAACAATAATTGACTGTCTATTTCTCGAACTTGCTATATTA
TTAAGCGCATTACTAAGTAATATAATCATTTATTTTGAATTTTGAGATTTGATATTAAAAAGTCATGCTATTTCAG

>Exon 2 VectorBase  (749 bp)

ACGATTATGGAGGTAATCGCTTTGGGAGAGGTGGAGGAAGAGGAAGAGGTAACCGAGGAGGAAGTTTTAGTGGTGGAAGA
GGTCGTGGGAGAAATGACCATGAATCTGCTGAAACTCCAAGTAATTTCTATAATTCTGCCAAGTATGAAAATAACGACGA
TGATTGGGAGGAAAACCCTAGAGACAGTGATCGTGTGAAGTCCAGCACCTTCAGAAGTGGCGGAAAAGGGTATGGTGGCT
ATGGTAGACAAGACCGTAATGTCACTTTTGATAATGATGATGATCAAGATAACCAGAGCGATGATGGTAACAGACAAAGA
GGAGGTGGAGGTAGAGGATCTGTGTCACATGACAGGAGAAATTTCGGTAACGATTGGGACAGTGGTAGTAAAAGTTACAG
CAGAGGTGTGGACAGATTTGGTGGTGGTGATGGTGGTGATTATCGAAAAGCTCGAAATGAAGATAGTCAAAGAGGTCGTG
GGCGTGGTAGGTGGGGAGGTAAAAATGATAATAATGATTCAGGTAATTATTATTCGAGAAAAAGAAACGAAGATAATTCA
CCTAGCAGGAGTGATGATGGAGATAGACGTAGAGGTGGTCAAAGAGGTGAAGGAACGGCAGAAGAGAAAGATGAGAAAAA
AAAAGAACGATATATTCCTCCGGATCCGACTGATGATGAAGAAGTAATGTTTGGCACTGGAATTACATCCGGAATAAATT
TTTCTAAATATGACAACATTAGAGTGAAG

>intron 2 VectorBase  (121 bp)

GTTAGATTTTATATAAACTATTTTCTAAATTTGTAGTTTTCAATTATAAAATCAAGTATTCAGTTTGAGAGAAAAAAAAC
TTGATTTTTTTTAATTTTTTTATTTATTTTTTTTTTTTTAG

>Exon 3  (186 bp)

GTTTCTGGTGAAAACTCCGTACCGCCGATAAATAAATTTGAAGAAGCCGGTCTAAGAGATCTAGTTTATAAAAATGTCAA

GAAATCCGGATACATCACTCCTACACCTATACAGAAGTATGCTTTGCCAACAATCATGATGGGTCGAGATTTAATGGCTT
GTGCCCAGACTGGGTCCGGCAAAACG

>Intron 3 VectorBase (112 bp)
GTAAGTTTACTTTCTGTTACTTTCACTTAATATGATTGCGGAAGAGAAAGAATTACTTCTCACTCAAAGTTTATAGAGAT
TTGTTTATTTTAAATCATTGTATTTATTCCAG

>Exon 4 (222 bp)
GCTGCGTTTCTCATCCCTATAATTCATACACTTTTGGAAAGTCCATCCGAGTTGATTATCACATCAACTTCTGTGGAACC
TCAAGCGGTTATCATGAGTCCAACAAGAGAACTTACTATTCAAATTTTTAATGAAGCTAGAAAGTTCGCCCATGGCTCCA
TTATCAAAGTGGCTATGGCATACGGCGGAACTGCTAGTTTCCACCAAGCTTCAGTCTTAATG

>Intron 4 VectorBase (114 bp)
GTAACTCTTATTTGATTTATTGTTATAGTTCAGTAGCTTTAATGGACCTATTCCTTCTAATAAACCACATTCGTACATAT
TAATCTACCTTTCTTATTTTCTTAAAATATCCAG

>Exon 5 (178 bp)
AGAGGCTGCCATATTCTAGTTGCCACTCCTGGCCGACTAAATGACTTTGTATCACGTGATAAAGTCAAGTTTTCTTCTGT
AAGATATTTTGTTTTGGATGAAGCTGACCGAATGTTGGATATGGGTTTCAAACCGGAAATTGAAAAAATGTTACTTCACG
AAAGTATGGTACCTGTGG

>Intron 5 present work (2758 bp)
TAAGTGAATTTATTAACTAATTTATTTTATTTTTTGTTGTTAGTCATATTTATTATTAATTTAAGAACGTTCAAAATAAT
TCTATAGAAAGCAAGAGTTTCTAATGATTGTATAAGTCTTCCATTTTCATGTTGCCTCTTCGACTTATAAATATCTTTTT
ATAACCTCGTTAAATCGTCTGTCATTGAGGTTTCATTTTATTCTACCCATACAATTCTTGTTTAGATTAAATTTTTTCCA
TTGTCGGGTCAAGATTCAGGCCTTACTGTGGCTAACAATTTATTAGTATACAGGGTGTTCATTTTAAGACGGCTCAAATG
CATTTCATGCTTAAACGGTTAATCTGAGCGAAAAATGGTAGAGGAAAAATTTGTTGCCCGTCACCTCTAGTTTCAGAATA
TGTGACTTGATTTTAGATTCCTACCGCATACGCGACTGACGCGCGACCAAAAGTGAGGGTTTGCAGTTGACCCCTATTTT
CATATACACCGTTTTAAAGAGTATAAAAAGTTACGAAAGAAACCGGAAACCACAATTCGATATCTTTAAAATTAAAGCTG
TAGGTCATTTCAAAATCTGGTATTTATATTTCAATTTTTTAACTTGTTATCAAAGAGAAAGAAGAGCACGCGATAGGTCA
TTTTTATGTATTTTTTCATCCCCTTTCTACTAAAAATATTGCCAATGCAGTAAAGCTAGTATTTTTCTTAATATCGGGAA
ATAAGTCGTTTTCCGAGTTATTTTTCTAAGAAAAATCAAATGTTAAGAGCTTTAAATGTTATCAAGCAGTACACAAAATG
GCTAAAAATGAATTTTACATCTGTTACAAACAATAAGGTTTAATAATTATTTGGTAATTTTGGGAGAGTTGCGTTTAATT
ACCAAAAAAATTGGAAACCACCTGCCTCATATTTAAGTAATCATTATCTTTCATGTAAAGGATCAAATTTATTCCTTAAT

TTATCAAATAATGCACTTTACACAACTTCTGAAGCATGTTCTGTGCCTTTGGAGGTTATCAAMTTTTYTMTSGATGGGGA
GGAGAWATYTTTTAGGGCTCAGWAMAAWRCTTCAGAAGTTGTGTGAARTGCATTWTTTGATAAATTAAGGAATAGGTTTG
ATCCTTTACATGAAAAGCCATTTTGTGTATGCTTGAGAACATCTAAAGCTCTTAACATTTGATTTTTCTTAGAAAAATCA
CCCGGAAATCGACTTATTTCACGATATTAAGAAAAATACTAGCTTTACTGCCTTGGCAATATTTTTAGTAGAAAAGGGAT
GAAAAAATACATAAAAATGACCTATTGCGTGCTCTTCTTTCTCTTTGATAACAAGTTAAAAAATTGAAATATGATTAGTA
AATACCAGATTTTAAAATGGCCTAGCACCTTTAATTTTGAAGATATCGAATTGCGGTTTCCGGTTTCTTTCATTACTTTT
TATACTCTTTAAACCTGTGTATATGAAAATGAGGGTCAACTGCGAACCCTCACTTTTGGTCGCGCGTCAGCCGTGTATGC
GGGCAGGAATCTAAAATCAAGTCACATATTCTGAAACTAGAGGTGATGGGCAACAGAATTTTTTCTCTACCATTTTTTGC
TCAGATTAACCGTTTAAGCATGAAATGCATTTGAGCCGTCTTAAAATAAACACCCTGTATAATGTTGCGACCATTTTTGG
CTACGCTTGTAAGAGCTGGTACTCTACATGTGGAAGTATACATGCTGGTACTCTACAAATTATGATACGTGTTTTTCAAA
TTACAGTTTGATGCTACAAGATGTACTAAAATAGATGGTCTACATGGCTACTAGATCTTGCTAGTATTAAGCATTTTATC
ATTGTTAAATAAAACGTTTTATTTTTTTATACATGTTTATGCAGCATCAATCCTGAAATAAGATAAATCTCGATTCATGA
CTTCATACCCACATGCCAAATATTAATCAAATACAATTTAATTTAAGGCATTTTTGGAATAAGGTTTTTGTTTCAATATT
TCCTTGCAAATGAAGGTTTTCTAATTCAAGTTAAGGTACATACAATAAGGAATGGATAATTGTATTACCCGGTTAAAACA
TCTAATGTTGCAAATGTTTATTTATGGGTAAATAAAATTCTATGTTAATCTGTAAATAGGCTTTAATTGAAAATAACGCT
TCTGTATTATTAATATTAGGCACTCATCCGTGAATCATGGTTCGCATGATTTTCTTATCCTATCTGGTTTCATATTTTCG
TCCACCTACAGAGGTGTACATCAGATCCCATGTCAAAAGTTTTGTCAAGAGATCAGTCATGTAATTGTGGCAAAAACTAG
ATATTTTTTAAGTTGAAATTGGAATTCAGATCAAAATTTTGTTTTCCATAACCCATCGGGAAAAAAACCCTAATCATAAT
ATATTAGTACATTAGTAATAAATATGTTTCTTTTCCACTCCATTTTGTATGTAGTGCAAATCTCGTGCAAAAAGTAAAAA
GTGTCATGAGTGTCGGTAGAGTTATTTCCCAATGGTAACTTAAAGCTTGTATACTTATTTCTTGGAATTTGAGTAATGAT
AGGTAATATTCACTCCAACTTATAATTATTAATTACTTAAGTTGGATCTTTTCTTTGGATTACACTTGGGTTCTTTTCAT
AATTTATAAACTTAACAAAATTTTACTTCATTGTTATTCTTATTTAACTTTAAAGGTGATTTTCATATTGCAGGTTATTA
TCACATAACACTTATATTATCTGCTATTCTCCTGCAGG

>Exon 6  (199 bp)

GGTCAACGACAAACAATGATGTTTTCGGCCACATTTCCTGAAGACATACAACGTTTAGCTCAAGATTATCTGAATAATTA
TCTCTTCATAGTTGTGGGCATTGTTGGAGGCGCTTGTACCGATGTACAACAAATATTTGAAGAGGTGGCACGCTTTGATA
AAAAAGAGAAATTAAAGGAGATTCTGTCCCAGACAGATG

>Intron 6 VectorBase  (1499 bp)

GTTAGTTTTAATATGAGCACTTAATAAAAGCAAAAAAAAAAACAAAAATCTAATTTAAAAGGACCTTGAATAAGCTCTCG
CAGTTTTTTGTTTTTGAAAATTAATCCTTTATTTTTAAAATTTGTGAAAAACAAATTATTGAAAAACATTTAATTTCATT
AAAACATTTATTTTATATATTGTCCCTCGCTAGCCACTACTTTTTACCATCTTTCTGGCAGCATTTGAATGACACTTCTT
TTGAGGAAATGCAAGAGTCGGTCCATTTTTGGATGGATTTTTTTTTGAATGGAGTGCTGCTCAGCCAAGCCATGTGCCAT

CGAGCGGAACAAGTAGTAATCAGAAGGGGCGGTGTCAGGTTTCCATTTAAGTGTTTCCAAATAGTTTTTCATCGGATGTG
CGACATGTGGCCAAGCGTTGCGATGTAGCAACATCACTTTGTCGGATCTCTGCTGGTATAGTGGCCGCTTTTCTTTCAAT
GCGTGGCTCAAACGCATTAATTGTAAACGATAGCGATTTTCCGTGATGGTTTCCTTCGGTTTTAGCAGTTCGTAATGCCG
AGTAAAATCTTAGAATCATGGATATTTGGCTTTGCTTTGATACATGCATGCTAGGTTTACCCCACGATCTTCTACGCCTA
GGGTTATCATATTCTATCTACTCTTCATCGCCAGTCACAATGCGATGCAAAAAACTTTTACTTTTCAGCCGTTGACGCAG
CAGTTCACACGTGATAAGACATCTCTCGTTTGGCACCCAATGTCCTTGCTTTTAAATCATTCCTACATGCTTTTAAACGT
TTCGAAACCGTTGTGTGATCAACTCCTAATGAACCTGCAAGTACAGCTAGCGTCTGACATGAGTCTTCGTGAAGTAATGC
TTCCAATTCTACACACTTAAATTCAAGTTTGGTTATAGGGATAAATTTTCACTATAAATATCCGTTTGAGCTACTCTTAG
CCCAAAAAAGTGGTTATAAGAAAATGGTCCAGTTTTGTTTAAACAGTAGATCAGAAAAACTCCTAAATCTCAGTAATATT
AATATTTCGTTTTGAGTACCGTCAAGGGATCTGACAGAATTTGACAAAGAAAGAAAAAATCCAAGTAGAAGTACTATACT
GCTTTATAGAATTTATTCCTCAATGAGTATCCCTTAATTCTAAGGTTTAATTCCTGTAAAATGTACTGCTTACTAATAGT
ATAGGTTTTTGAAATTCGATGTTCGAAAAAATAAGTAACTGAGAATTGTAAGGAAAAGACGTGCGTTTTCACGCCTCCAC
AAAAAAAAAAAAAAAAACATGCAAGAGGGTTTTTTTGTTTTATTGACACGTGCTCAACTATAAAGCTTTCACTTAGAAAA
TTTTTTATTTTCTCATAAGTTTGAAGAAAATTTATTTAAAACTGTGAGTATGTACAATAGTTTGAGCAATTTGTATGTAT
TTCACGTAATTTTGTTTTAAAGATAACATCTAAAAATAATAAATTGTATTTATGAATAG

>Exon 7  (198 bp)

ATAAGGAAAAAACGTTGGTGTTTGTGGAGCAGAAACGAACAGCCGATTTCATTGCGGCATTTCTTTCGGAAAATAATTTT
CCAACTACATCAATTCATGGCGATAGACTTCAGAGTCAGCGAGAATCTGCGCTTAATGATTTTAAAACTGGGCGTATGCC
AACTCTTGTGGCAACAGCGGTCGCAGCAAGAGGTTTAG

>Intron 7 VectorBase  (1370 bp)

GTAATAAAATATTTGTTTATTTTTCTTAAATGTAATTGTTTTAATGGATGAACAATATTAAATGAAATCCACTTTTTGTT
AATAAATCGAATTATTAGACTCCTGTGTTTAGGGATTAAAACGTTTTTGCTGTTTAACAAAACTATTGAACTTCATGTGC
TTGTAACTTTAAAGTTACGTATTCACAACTGTTTCCGAAGTATACATCCTTCACCAGTTTTTGGTAATTTTGCCATTTAG
TTAAATGCAGCTTTGTGTTAATTGAAACATTAGTGAAGTATAACTTGTGCCCTCAAGTTTCTGAAATAGAGACTGACTGA
TTGAGGTCACATCAATTCAATCCACATGCTCCACTGAGATAGTCATAACAGGATTTATCTAAAATTTGCTGTGAACAGTC
ACGTAGTTAGATCTGTAGAGCAAAAAAAAAAAAAATGTCTACGATCTTTTCTGTGAGGGGCTTGCTTTATTTTATGATCAA
TGGTTTTCTTCAAGATGAACATTGTGGGAAATAGTGTTAGGTCAATGTGCCCTTCAAAATCAGTTTTGAAAAACCATCAC
AGTAGTTATAATAGATTAATTTGGAAATCCCCTGAATAAAAGAAATGAAAATGTTTTCTCAGAGCAATGGCAGTGTGTGT
TTTCAATATTTATGTCTAAAAGTAAAAGTCGCAGAATGCCTCAAACTTAGTATTCGGCTTAGGTATTCTGTTTTCACTTC
TGATAACCAAGGGTATTAATTACTACAACGAAATTGCGAATCTATTGATGCGAGTGTTACTTTTGTCTCAAAAGTTACTT
CTTCAAAATGTTACACTGACGAATAAGATTATTGTTAGACTGGATATGAAATTAATAGATATTTTCTAAATTCTGAATTT
ATTTCGAAATCGGACTTAGCTGTTTTTATTTAACATGGAATAACTTTATGTAATTACCATATTTCCCAAAATCTGTTAGT

TAACATTAACTAACAGATTTTTTAAGATACGAGTATTTAAAGTTTTTTTGTTTAGCTCTTTTCAGAAAATTCTGTTTGAA
GTTTGAAAAGGACATGAATATGCATCACAAATACGCTTTTCAGTTAGGCCACCCCGCCGAAATAATCATGGCTTTTCAAT
TAATACATCTTTTATTTTTCTTTCAAAAATGTCAAATTTATTTAAGTTTAATTAAAATAAATATTAATGTTTTATATCTC
ATAAAAAATTTGCTTTTCTCACTTATCCTCTTTTTCAAAGTAACTGATTTAATTTTAGAAAATATGAATTTTTCATGACC
TAATTTCGCGGAGTAAATTATCCATGTCAAAGCTGCTTGTTTCTAGTTTACAAAGTGCTGGTTAAAGCTAATTTATTTCT
GTAATTACAG

>Exon 8  (173 bp)

ATATAAAAAACGTGAGCCATGTTATTAACTACGATTTACCAAGTACAGTTGATGAGTATGTGCATCGAATTGGTCGGACT
GGTCGAGTTGGTAATCGTGGCAAAGCTACATCCTTTTATGATCCTAATCAAGATTCAGCAATTGCAGATCAACTAGTCAC
CATATTAAATCAA

>Intron 8 VectorBase  (91 bp)

GTATTAAATTAATTTTTGTACAATCTGTGTCCAAGTAGCTTTCTTCAAACAATATAGTTTAATAACTTAATGATTATATT
TTTATTTATAG

>Exon 9  (126 bp)

GCAAATCAGCCAATACCTGACTTTTTGACATCATCGATGAGTGGAGCAAAATCAGAATCACGTAGTTTCGGCAATGATAA
TATTAATCAAGGCTTTGGTGGCAGAGACGTCCGAAGAGGACCTTCG

>intron 9 VectorBase  (464 bp)

GTTGGTAGACCTTTTGTTTTATTACTGCATTATTATTATAATTTGTTAGATGCTTCATTCAGATTCATTGCTTCTTATTC
GTTACTCACAATTTGCTTCTTAGCATTTTTGTTAGTGTAATAAGGATGGTTTAAATATAAATAAGCAAGGAATTTTTTTG
CTTCAAGATTAGTTCATTGTATAAAGTTTTGCAGTCGCAAAATTATGGAAAAAATATAAATATTTATTTGGTTGCGACAT
TAAGCTTATTATTACTTTGATAAAAATCCAGACTAAGTGCACTGTTCGCGTTTAATTTTTGCCTGCTCTTCAGAAAAACA
GATTTTAGTTTTTGTTAACTTGCTACTGTTTAGAGGGCGGCCAGTATTATTGTTGAACTAAAACCACTTGTCCATTTCAT
AAATTTTTATGATTGTCCAGTAGGAACTTTGATAATTCTCTTATACATTCTTTTCTCTTTGCAG

>Exon 10  (57 bp)

CAAAGTTACCAAGCAGATCGATCTCCTCACGTCCAAGAAGATTATGAAGAATGGTGA

>3' mapped  (982 bp)

AAAGTGGAATTGATAAATAGATTAGCTCTGAATGCGGAGTTGAATTTTGAAATATAATATGTATTTTCTACTTATCTGAC
TTTGTTAAAAAGATACAAAAACTTTTCTTTAAAGTACAAAAAAAAAAGAGTAAAACTTTTAAATATTTAGAAGAGACTTT

TTAGAGCATTTATTAGTTTCATGTTATTTTTTACATACATTTTTTTTTTTCAAGTAAAAAGTACGCGTGGTGCAGTGTGT

TTTCGTTTAAATATTAAAAAATATGACTTTATATTTAGTCTTCAATGACGTGATAATGGTTCTTCTGTTAGATGCAAAAT

CTCGTTAAAAAACAAAAAACAAACACAAACAAAAGTGCCTTCAAATGTGATAAACTAGTTCCTTCATTAATAGAAATGTT

GCTTTTTTTAAACAAAATAAGCATAAGAGATATTATTTTGAATAAAAATAAGAAATCTTCTATTATTAAGATAGATAAAT

CTCCTAAGTGATATTTAGTTACCTAGAAACAACAATCGTATTTTTTTCACAAGGCAAAACATTATTTTTCTATTAAAATG

AAAATATTTCAAAATAATTTTTTTTTTTTTAAGTGAGTTTATTTGTTGCATAGCTGAACTACTGTTTGTATTCCCAAGTA

CTTAGTACTTTCAGTTATCGAATATGTTTATCTTCCTTCTAGATTTTATTTAGGAAGATTTTTTCTATTTCATTAATTAC

AGTGCATTATAAAAATGGAAAAGAAAAAAGAAAACATTTAATTTTTATAATATTAATTTGTTTTTATTATATTTTCTAAA

TAAAATATTTTCTTTTCTATTTATCAGTTATTGTTCTCGTAGATAAATTTAAGATTCTCGCTTCGATCTTCGCGAAAAGC

GATTCTGGCCAGTTCTGACCAAAAAAACCAGTGACCCAAACAACTGTCCGACATGGTGAATCACTTTTCCAGGAAGCCAG

TCGTGCACTACTAAATTCACAA

Martins et al. Figure S5

```
                                       *        20         *        40         *        60         *        80         *       100
D._melanogaster_zpg     : -MYAAVKPLSKYLQFKSVHIYDAIFTLHSKVTVALLIACTFLLSSKQYFGDPIQCFGDK-DM--DYVHAFCWIYGAYVSDNVTVTPLRNGAAQCRPDAVS :  96
D._melonogaster_inx2    : -MFDVFGSVKGLLKIDQVCIDNNVFRMHYKATVIILIAFSLLVTSRQYIGDPIDCIVDEIPL--GVMDTYCWIYSTFTVPERLTGIT--GRDVVQPGVGS :  95
S._americana_inx2       : -MFDVFGSVKGLLKLDSVCIDNNLFRLHYKATVIILIAFSLLVTSRQYIGDPIDCIVDEIPL--AVMDTYCWIYSTFTIPNRLNGKI--GLEVAHPGVGA :  95
S._gregaria_inx2        : -MFDVFGSVKGLLKLDSVCIDNNLFRLHYKATVIILIAFSLLATSRQYIGDPIDCIVDEIPL--AVMDTYCWIYSTFTIPNRLNGKI--GLEVAHPGVGA :  95
N._vitripennis_inx2     : -MFDVFGSVKGLLKLDTVCIDNNVFRLHYKATFIILVAFSLLVTSRQYIGDPIDCIVDEIPL--HVMDTYCWIYSTFTIPDR-NGVV--GKDIVQPGVAS :  94
A._melifera_inx2        : -MFDVFGSVKGLLKLDSVCIDNNVFRLHYKATVIILIAFSLLVTSRQYIGDPIDCIVDEIPL--HVMDTYCWIYSTFTIPDR-TGVV--GKDMVQPGVAS :  94
T._castaneum_inx2       : -MFDVFGSVKGLLKIDVVCIDNNVFRLHYKATVIILIAFSLLVTSRQYIGDPIDCIVDEIPL--NVMDTYCWIYSTFTIPNRLTGRV--GLDIVQPGVAS :  95
A._aegypti_inx4         : -MLEITKSLRDIIVPKSFDSTNTVWRLHSRITVYMLVFETILLSARSYFGEPIECISSAAPTVRASLHSFCWTLGTYISRDPNFVEA--SWDIIEIGTHM :  97
A._gambiae_inx4         : -MLEFVRPLQSIIQIKQVNSTDLVWRLHCRVTVFLLILASLLLSARQYFGNPIDCVIGSGTVSSSTMNEFCWIMGTYISNDPNFVLD--STDLVKINAKI :  97
R._prolixus_zpgA        : -MFDVFGNLKGLLRIDSVCIDNNVFRLHYKATVIILVAFSLLVTSRQYIGDPIDCLVDEIPL--NVMDTYCWIYSTFTIPNRLGGTV--GKDVVQPGVAS :  95
R._prolixus_zpgB        : MLLEIFTFLKLIFIEKAVNIDNLVFKLHYRITALLLFIFSVLVTARQYFGDPIDCLVQGVPE--KVMDSYCWISSTFTVRYLPHQQI--GVHMVQPGVKS :  96

                                       *       120         *       140         *       160         *       180         *       200
D._melanogaster_zpg     : KVVPPENRNYITYYQWVVLVLLLESFVFYMPAFLWKIWEGGRLKHLCDDFHKMAVCKDKSRTHLRVLVNYFSSDYKETHFRYFVSYVFCEILNLSISILN : 196
D._melonogaster_inx2    : HVEGEDEVKYHKYYQWVCFVLFFQAILFYVPRYLWKSWEGGRLKMLVMDLNSPIVNDECKNDRKKIILVDYFIGNLNRHNF-YAFRFFVCEALNFVNVIGQ : 194
S._americana_inx2       : HVAGKDEVKYHKYYQWVCFVLFFQAILFYIPRYLWKTWEGGRIKMLVLDLNSPVVNEQSKADRKKLLVDYFATNLHTQNF-YAYRFFICEALNFVNVVGQ : 194
S._gregaria_inx2        : HVAGKDEVKYHKYYQWVCFVLFFQAILFYIPRYLWKTWEGGRIKMLVLDLNSPVVNEQSKADRKKLLVDYFVTNLHTQNF-YAYRFFICEALNFVNVVGQ : 194
N._vitripennis_inx2     : HVDGEDDIKYHKYYQWVCFTLFFQAILFYIPRYLWKTWEGGRIKMLVLDLNCPVVSEDCKTDRRKLLVDYFATNLHSQNF-YAFRFFLCEVLNFINVVGQ : 193
A._melifera_inx2        : HVEGEDEIKYHKYYQWVCFTLFFQAILFYVPRYLWKTWEGGRIKMLVLDLNCPVMSDEFKSERRKLLVEYFATNWHTQNF-YAFRFFLCEVLNFVNVVGQ : 193
T._castaneum_inx2       : HLDGTDEVKYHKYYQWVCFALFFQAMLFYVPRYLWKTWEGGRIKMLVLDLNYPIVSEDCKTDRKRLLVDYFITNLHMQNF-YAFRFFICEVLNFVNVVGQ : 194
A._aegypti_inx4         : GHIPKEERLYQKYYQWVPFLLAIQAFLFSFPKHLWRFCERGRLETLCHNLTSILSPGAWTRKRKALTLLYLTQESRKGHNKYALIFIGCEILNFFIVLLN : 197
A._gambiae_inx4         : GHIPESERSYQKYYQWVVFILALQACMFSVPNFLWKAWEAGRLQSLCDGLTTPIVPDHWEKTRKKQLITYLSADFPRLHRTYLLRYCFCTLLNFCNVLLN : 197
R._prolixus_zpgA        : HVEGEDEVKYHKYYQWVCFVLFFQAILFYIPRYLWKTWEGGRIKMLVLDLNCPVISEDCKEDRKKLLVDYFTSNLHTQNF-YAFRFFICEILNFVNVLAQ : 194
R._prolixus_zpgB        : YIEGEDTVKYHTYYQWVCFVLFFQAVLFYTPHYLWKNWEGGKIKELVEDLKTPFSKEKNIKQRKKIIINYFKNSKGQHNV-YIYHFLLCELLNLCNVFAQ : 195

                                       *       220         *       240         *       260         *       280         *       300
D._melanogaster_zpg     : FLLLDVFFGGFWGRYRNALISLYNGDYNQWNIITMAVFPKCAKCEMYKGGPSGSSNIYDYLCLLPLNILNEKIFAFLWIWFILVAMLISLKFLYRLATVL : 296
D._melonogaster_inx2    : IYFVDFFLDGEFSTYGSDVLKFTELEPDERIDPMARVFPKVTKCTFHKYGPSGSVQTHDGLCVLPLNIVNEKIYVFLWFWFIILSIMSGISLIYRIAVVA : 294
S._americana_inx2       : IYFMDLFLDGEFTTYGSDVVRFTEMEPEERSDPMSRVFPKVTKCTFHKYGPSGSVQTFDGLCVLPLNIVNEKIYVFLWFWFVILSVLTGIGLVYRLATAM : 294
S._gregaria_inx2        : IYFMDLFLDGEFTTYGSDVVRFTEMEPEERSDPMSRVFPKVTKCTFHKYGPSGSVQTFDGLCVLPLNIVNEKIYVFLWFWFVILSVLTGIGLVYRLATAM : 294
N._vitripennis_inx2     : IYFMDFFLDGEFTTYGSDVVKFTEMEPEERVDPMSRVFPKVTKCTFHKYGASGTVQKFDGLCVLPLNIVNEKIYVFLWFWFIILSVLSGLSLAYRAAVVA : 293
A._melifera_inx2        : IYFMDFFLDGEFTTYGSDVVKFTEMEPEERIDPMSRVFPKVTKCTFHKYGASGTVQKFDGLCVLPLNIVNEKIYVFLWFWFIILSVLSGLTLAYRAAVIA : 293
T._castaneum_inx2       : IFFMDYFLDGEFSTYGRDVLSFTEMEPEEREDPMSRVFPKVTKCTFHKYGPSGSVQKFDGLCVLPLNIVNEKIYVFLWFWFVFLSVLSGLSLIYRLVVIF : 294
A._aegypti_inx4         : MFLMNFLFGGFWASYQPAIQALLSLDMNAWTSYNSLVFPKLAKCDFSYIGPSGSKQNFDALCLLPQNIVNEKIFAFLWLWFIVLAVVSGVQLCYRLAQLS : 297
A._gambiae_inx4         : IFLVNVIFSGFWSNYHPAVKALLSFDFPSWNRYNSQVFPKIAKCDFHFVGPSGSKQNRDGLCLLPLNVVNEKIFAFIWLWFLGLLVISMLNLLFWIVVLC : 297
R._prolixus_zpgA        : IYFMDFFLDGEFTTYGSDVVRFTEMEPEDREDPMARVFPKVTKCTFYKYGPSGTVQRFDGLCVLPLNIVNEKIYVFLWFWFILLTLLSGLSLIYRAAVVA : 294
R._prolixus_zpgB        : IYLLDVFLDGGFINYGYHLIGGYDDSATMNLDPMASLFPKVTKCTFKKFGPSGTLQKFDGLCILPVNILNEKIYIILWFWFYILAVVTSLWLLFELIVLA : 295

                                       *       320         *       340         *       360         *       380         *       400
D._melanogaster_zpg     : YPGMRLQLLRARARFMPKKHLQVALRNCSFGDWFVLMRVGNNISPELFRKLLEELYE------------AQS---------------------------- : 356
D._melonogaster_inx2    : GPKLRHLLLRARSRLAESEEVELVANKCNIGDWFLLYQLGKNIDPLIYKEVISDLSR------------EMSGDEHSAHKRPFDA--------------- : 367
S._americana_inx2       : GPQMRMYLLRARSRLAPQDQIETISNKCQIGDWFVLYQLGKNIDPLIYKELVADLAK------------KLEGKEIV----------------------- : 359
S._gregaria_inx2        : GPQMRMYLLRARSRLAPQDQIETISNKCQIGDWFVLYQLGKNIDPLIYKELVADLAK------------KLEGKEIV----------------------- : 359
N._vitripennis_inx2     : GPKLRFILLRARSRLSHQDQIEVISNRCQIGDWFILYQLGKNIDPLIYKTLIADLAK------------KFEGKENV----------------------- : 358
A._melifera_inx2        : GPKLRLVLLRARSRLSKPEHINTIAEMCMIGDWFVLYQLGKNIDPVVYQQLVVDLAT------------KLQGKESV----------------------- : 358
T._castaneum_inx2       : MPKVRLYLLRGKCKIAPQKEVEIINTRCEIGDWYVLYQMGKNIDPLIFREIISDLSK------------KLEGKETV----------------------- : 359
A._aegypti_inx4         : CRSVRFQLLFSLLDPISYHRLKRVVREANIGYWFLLYQMARNINKGVMREIIRDLSRIDQ--------ELNMSKSNVNELQLEVEAEDELEEIEDDEATV : 389
A._gambiae_inx4         : SKGFRLWLLTAPLYPIRTSYVARALDGQGVGQWFLLYQLCRNLNPIVGRELVQSVSKAKGHNGHKTFRMPKGGEPDFYTDPEGYDEEGV----------- : 386
R._prolixus_zpgA        : GPQIRLYLLRARSRLSPQNHIEIITNKCQIGDWFVLYQLGKNMDAIIFKELLADLAK------------KLEGKETV----------------------- : 359
R._prolixus_zpgB        : SSKLRFSLLKARAKIVNTNTVNSILKKFKIGDWFLLYQISKNMEILTLKSFLEELKR------------KLNNGKPK----------------------- : 360
```

Martins et al. Figure S6.

```
                          *        20         *        40         *        60         *        80         *       100
D._melanogaster : MVADNIDAGVA----------IAVADQSSS-PVG------DKVE-------IPAGNYILVGVDIDTTGRRLMDEIVQIAAYTPTDHFEQYIMPYMNLNPA :  76
R._prolixus     : ----MANKEGP----------------------------------------------VYLSWDIDTTGRRLIDEICHIAVHDVYSSFEQYVMPFRDIDYS :  50
A._melifera     : MVSSTIMENGK-----------SNPSRKPS-AVG-----------------IGPGNYRLVGWDMDTTGKKVIDEICQIAGYTPSSSYSQYVMPYKDLNPP :  71
T._castaneum    : MVQNDSSNSGD---------EGSPPVAAPNPAKG-----------------IPLGKYRLVGWGVDTTGRRLIDEICQIAAYSPSSQFSQYIMPFSDLNPS :  74
A._aegypti      : MVTGGIDESKTEGEFDQQQLDESCSDESDTGPAGALSSKFDRTDVSINTNPLPYGKYTLIGIDIDTTGRRLIDEIVQISAYTPEHQYSQIYMPLMNLNPA : 100
A._gambiae      : MVV-GMDESKV-GHDEEQHYEDSSFDECDT-PAGVLSSKFDKTDLSINTNPLPHGKYTLIGLDIDTTGRRLIDEIVQISAFTPEKEYAQYIMPLMNLNPA :  97

                          *       120         *       140         *       160         *       180         *       200
D._melanogaster : ARQRHQVRVISIGFYRMLKSMQTYKIIKSKSEIAALKDFINWLEQLKTKAGPSSDGIVLIYHEERKFIPYMILESLKKYGLLERFTASVKSFANSINLAQ : 176
R._prolixus     : ARKRYLLKTVNIGFYRVLRETASGKILKTKSEISALTDFVVWLEQRKITEGRD---LILMYYETYTPAPFILIESLRKYNLLPRFKEAIKGFANSFSYIK : 147
A._melifera     : AMKRHNMKIVTIGKFRVLKDNKTNKALKTKSEVSALTDFITWLS--SIRGDAA-DGIILVYHEPRKVIPAMLIESLKKYNLLERFKQIVKGFANGFNIAE : 168
T._castaneum    : YRRKHSIRVVNTGRYRMLKDMRSNKFVKTKSDVAALTDFLEWLE--KVQGDDAHDGIILIYHEIRKASPGMLLEVLRRYNLLERFAKIVKGFANGFNIAQ : 172
A._aegypti      : ARQRHQVRVITVGFFRMLKSMQTYRVVKTKTEISALNEFINWLEE-RYREDEGSEGIVLIYHEQRKFVPYMLIEAVKKYNLLDRFKSIVKSFANGFKLAE : 199
A._gambiae      : ARQRHQVRVITVGFFRMLKSMQTYRVMKTKMEVAALNEFLDWLEE-RLREDEGSEGIVLVYHEQRKFVPYMVIEALKKYKLLDRFSESVKSFVNGFKLAE : 196

                          *       220         *       240         *       260         *       280         *       300
D._melanogaster : ASIGDANIKNYSLRKLSKILSTTKEED----AACSASTSGSGSGIGSGSSMVS-------DSVSISPRDSTVTNGDDKQSSKNAVQGKRELFDGNASVRA : 265
R._prolixus     : SKCEKI-MQTYSVKNIAKVLLNKEYKS-----HDACERARILYEIVAHISVGE------------------SQDGE------------------------ : 199
A._melifera     : VKCANT-VRAYTLRTLSQSLFNQETQL-----GNAKDRACLALQIVQHLSSLE------------------VTKGNEAD-------------------- : 223
T._castaneum    : AKCAKT-TKSFSLRVMAKVLLNREDED----FSSAVDRARVSYEIAAHLAQGE------------------RQDLDEKK--------------------- : 228
A._aegypti      : EKCSKT-IKYFSIRQLAKIVLDESEAK-DGFEGNAAYRARLAYQLARRLSKEDDEKPPTNDSTQVAAECEETQSDDEAETEEGAVGGSNTAEDEAKSQES : 297
A._gambiae      : EKCTKT-IKYLTIRQLAEIVLDEKAGSRDGFEGNAAYRARMAFEISRCLATAE----PKSNAVNVPASSPTESEPNDSATSEEVTNSSSTAEDAVKDEGD : 291

                          *       320         *       340         *       360         *       380         *       400
D._melanogaster : KLAFDVAL----------------------------QLSNSDGKPEPKSSEALENMFNAIRPFAKLVVSDVLEIDIQIENLERQNSFRPVFLNYFKTTLY : 337
R._prolixus     : -----------------------------------------------------EDIAAALESYMSPVEEIEDGIAAMKLTIAKQNSLKPIFGQFIRMDKK : 246
A._melifera     : --------------------------------------------GSGDSDSAMKNTVEFIREFVQPVEIEEQEYAELKIVFERQNSLRPIFGVLFRVNRR : 279
T._castaneum    : --------------------------------------------PTGTE----PEIINFVCPEVNPISAEEDEIAQFKVLIERQNTFRPVFGALLRASRP : 280
A._aegypti      : GAATVNAENNGGPQGQNDSDDDRHQTDVDAVTGQVTTIPEPQPRSRPTTEAEREKMCSVLCDLASPITSEISELDEQEKILIRQNSLRPVFLQYFKTTIY : 397
A._gambiae      : KPSEENAE---------QSDDDAQKQDEKKEEAPAASLNSSA--SRPLTEEDREKMCAVLCEYASPISTEISELDEQEKILVRQNSLRPVFLLYFKTTIY : 380

                          *       420         *       440         *       460         *       480         *       500
D._melanogaster : HRVRAVKFRIVLAENGFDLNTLSAIWAEKNIEGIDIALQ-SI-GRLKSKDKAELLELLDSYF-DPKKTTVKPVVK----------------------GNS : 412
R._prolixus     : RRQ-ALILRRHLINSDIEYEMLKEVWT--RKEDMKVFLKKNIKDEISDDDINDLDFMINKHLGGGGGQTEKTKVS------------------------- : 318
A._melifera     : ERQHASPLRRLLAEAGIEYSQLQEAWSNGKKEGIDKLIKEKL-TATDEKKIEDVLIVLEGHF-DPDK---KSKLK------------------------- : 349
T._castaneum    : ERQHATHLRRLLAENNINYDKLKCAYEDGAKDGIDKVLKSEV-ANAKETELSELLDILDCFF-DPEKKAVQPKPR------------------------- : 353
A._aegypti      : HRVKAVTYRRVLAEYGHDFASLQCVWNDSKREGIEAVVN-KI-TELKEEERTELVELLDCHY-DPEKQQLKPVVKRVRGRSSR-GRLFFGNKSNNVSGNG : 493
A._gambiae      : HRVKAVTYRRVLAETGHDYESLRQVWQEKKREGIEAIIN-EI-AELKEDERTELNELLDCHY-DPDKQAFKPIVKRMKRRPSRAGQPFFSNKNGNVPQNQ : 477

                          *       520         *       540         *       560         *       580         *       600
D._melanogaster : NNNNNYRRRNRRG--GRQSVKDARPSSSPS-----------ASTEFGAGGDKSRSVSSLPDSTTKTPSPNKP-RMHRKRNSRQSLGATPNGLKVAAEISS : 498
R._prolixus     : -----IGRNSPKKNISRQAFRGNQRKPNR------------KFTDHSKSGKINPTNNGNPTNTTVATAKSSSNKSAGEKSNKTESTSEIRTDGESTTVPT : 401
A._melifera     : -----ISRDKS------------------------------QIKMKNSNINDDRENNNKCDSGTETPDSTTPNSPLKIK--------------------- : 393
T._castaneum    : -----YYQNNN---------------------------------NRKPRGTRKSFNKDRNSESTSSPNASSDNATEKTG--------------------- : 394
A._aegypti      : NGNGGNYNAANNGNVGNGNAKDQRKAVHGGRNNYHNNSN--ADMNNNQGGGKNNYNNNSGNNQPPSPNGGANGKKFRPRQMRRRR-------NQAGGR-Q : 583
A._gambiae      : NGNNQMAKENRKPFNGYQNNQKNHNNHHNGNNNNNNNNNMVNKHQNQQNQQQQQYYNGPSSPESGMRNGGSPGKRFFQRNSRRRRGGNGQNQNQNGGQNQ : 577

                          *       620         *
D._melanogaster : SGVSELNNSAPPAVTISPVVAQPSPTPVAITASN--  : 532
R._prolixus     : TTVTETLDSI-----------NTSSTPNSVDNKNEP  : 426
A._melifera     : ---------------------SESSENSTFIAEE--  : 406
T._castaneum    : ---------------------TESEKEKTVEVQ---  : 406
A._aegypti      : NAM------------------NQQQQPQMIHANA--  : 599
A._gambiae      : QPMKN-HNGNHHAYSNSNRHNNQFQQPMQVHA----  : 608
```
